# Comparing rapid rule-learning strategies in humans and monkeys

**DOI:** 10.1101/2023.01.10.523416

**Authors:** Vishwa Goudar, Jeong-Woo Kim, Yue Liu, Adam J. O. Dede, Michael J. Jutras, Ivan Skelin, Michael Ruvalcaba, William Chang, Adrienne L. Fairhall, Jack J. Lin, Robert T. Knight, Elizabeth A. Buffalo, Xiao-Jing Wang

**Affiliations:** Center for Neural Science, New York University, NY, USA; Department of Physiology and Biophysics, University of Washington, Seattle, WA, USA; Washington Primate Research Center, University of Washington, Seattle, WA, USA; Department of Neurology, University of California, Davis, Davis, CA, USA; The Center for Mind and Brain, University of California, Davis, Davis, CA, USA; Department of Psychology, University of California Berkeley, Berkeley, CA, USA; Helen Wills Neuroscience Institute, University of California Berkeley, Berkeley, CA, USA

## Abstract

Inter-species comparisons are key to deriving an understanding of the behavioral and neural correlates of human cognition from animal models. We perform a detailed comparison of macaque monkey and human strategies on an analogue of the Wisconsin Card Sort Test, a widely studied and applied multi-attribute measure of cognitive function, wherein performance requires the inference of a changing rule given ambiguous feedback. We found that well-trained monkeys rapidly infer rules but are three times slower than humans. Model fits to their choices revealed hidden states akin to feature-based attention in both species, and decision processes that resembled a Win-stay lose-shift strategy with key differences. Monkeys and humans test multiple rule hypotheses over a series of rule-search trials and perform inference-like computations to exclude candidates. An attention-set based learning stage categorization revealed that perseveration, random exploration and poor sensitivity to negative feedback explain the under-performance in monkeys.

## Introduction

Animal models are essential for mechanistic investigations of the circuit underpinnings of complex computation. New frontiers in the training of non-human animals to perform computationally challenging tasks while simultaneously recording from large neural populations across several brain regions promise rapid advances in our understanding of the neural substrates of cognition. However, our ability to extrapolate any findings to an understanding of human cognition relies on an overlap between the computational and neurocognitive means used to carry out complex tasks across species [1, 2]. The prefrontal cortex, which plays an essential role in higher cognitive functions [3], is disproportionately enlarged in humans compared to macaque monkeys [4, 5]. It has been argued that the resulting increase in the number of neurons may underlie superior human cognitive abilities [6, 7]. Thus, interpretation of findings from animals demands systematic and rigorous comparisons between cognitive computations in humans and non-human animals.

Towards this end, we compared the behavioral strategies employed by macaque monkey and humans on the same task: a variant of the Wisconsin Card Sorting Test (WCST), widely used to evaluate the cognitive functions involved in abstract thinking, rule search, cognitive set shifting and the effective use of feedback [8, 9]. The WCST and its variants have long been employed in the study of prefrontal function and dysfunction [10, 11, 12, 13, 14], lending support to the presence of abstract thinking and computation in the monkey brain [15]. In the task, subjects must match or select items composed of multiple visual features based on an uncued or hidden rule. Feedback at the end of each trial indicates whether the response was correct, but does not unambiguously reveal the rule identity. Instead it helps narrow down the rule identity, which must then be inferred from the collective outcome of multiple trials. Critically, the hidden rule changes in an uncued manner across trial blocks, requiring detection of and adaptation to these uncued rule switches based solely on positive or negative feedback.

We implemented a version of the task in which each card has three feature dimensions (color, shape and texture), each of which can take one of four values, defining twelve possible “rules” (Fig. 1a). What strategy might one use to identify the current rule from the outcome of choosing an object in each trial? It is clearly extremely inefficient to simply learn the value of 64 individual object-reward associations; each object is a conjunction of multiple visual features, and learning the value of one object does not generalize to any of the other objects. Learning the value of features to identify the rule is far more efficient as it allows generalization. In reinforcement learning, the problem of attributing binary feedback in situations when there are many features (high-dimensional environments), referred to as the “curse-of-dimensionality”, is effectively resolved through such abstract reasoning [16, 17, 18]. However, a rule-based strategy still poses challenges of cognitive capacity. Simultaneously tracking and updating the value of twelve features can impose prohibitive working-memory demands and be computationally daunting. On the other hand, selectively attending to and evaluating a single feature at a time is inefficient as it discards relevant feedback regarding other features. The strategy that humans or monkeys use to address this tradeoff between computational complexity and information efficiency remains to be elucidated.

**Figure 1:**
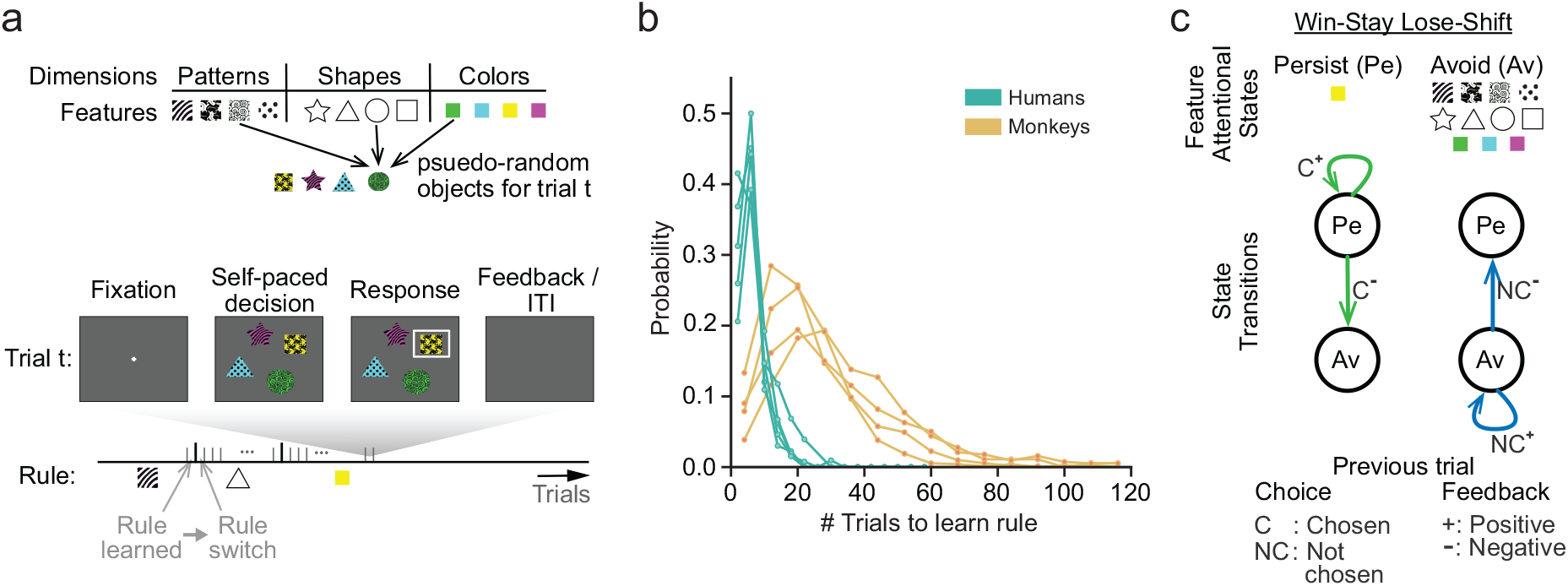
Monkeys rapidly learn rules in the WCST but are slower than humans. **a**. WCST task structure. Each trial is composed of fixation, decision, response, feedback and ITI epochs. After fixation, the subject is presented with 4 objects that are pseudo-randomly composed of 3 features - a pattern, shape and color. The features composing each object are mutually exclusive with respect to other objects. Each block of continuous trials is governed by a rule (one of the 12 features). The subject receives positive feedback only for choosing the object with that feature. The identity of the rule is hidden and must be discovered. An uncued rule switch to a random new feature occurs when the subject demonstrates they have learned the current rule. **b**. Distribution of trials-to-learning-criteria in 4 monkey (brown) and 5 human (green) subjects. All subjects rapidly learn the rule, but on average, monkeys are over 4 times slower than humans. **c**. Decision process for the Win-Stay Lose-Shift learning strategy in two-armed bandit problems. The decision to choose an arm can be in one of 2 states: persist when it is chosen and avoid when it is not. The decision to choose an arm stays in the persist state as long as positive feedback is received (win-stay) and switches to the avoid state otherwise. It then stays in the avoid state as long as positive feedback is received, and switches to the persist state when negative feedback is received (lose-shift).

To analyze how individuals handle the cognitive complexity and demands of this task, we developed a detailed behavioral model that allowed us to compare the strategy and performance of humans and trained macaque monkeys on the same WCST analogue. This allowed us to identify differences in the types of errors in the two species, revealing the underlying cognitive differences between them. Indeed, differences in the types and prevalence of errors on the WCST, demonstrated via similar comparisons between healthy and patient populations, has played an essential role in establishing behavioral markers of various forms of cognitive dysfunction [10, 19, 20, 21]. We found that while monkeys rapidly identified new rules and learned several rules over an individual session, the rule learning in monkeys was 3-4 times slower than humans. To understand this difference, we fit a behavioral model that predicts upcoming choices based on choices and their outcomes on previous trials. We modeled the transformation of the trial history to an upcoming choice through inferred hidden behavioral states. While the model is not constrained to do so, the best-fit models for all subjects of both species developed hidden states that aligned with a feature-based attention strategy wherein some visual features are selectively examined over others while making a choice and attributing feedback. The decision process governing each species’ rule search strategy is characterized by the statistics of the inferred transitions between these states. This strategy bore a striking resemblance to the Win-Stay Lose-Shift strategy but with a few important differences. First, our model revealed that both species often explore more than one features at a time. Second, both species perform inference-like computations – the model reported changes in the attentional state towards one feature based on the outcome of choosing another. We developed a novel approach to identify distinct stages of rule-learning. Analysis of these stages revealed three key reasons for slower monkey rule-learning performance: (*i*) following a rule switch, monkeys perseverate on the previous rule more than humans; (*ii*) monkeys often make seemingly random choices that do not involve any of the features under exploration, even after finding the rule, which delays the expression of the learnt rule; and (*iii*) poorer attention to negative feedback in monkeys particularly when they simultaneously explore the rule and non-rule features poses a credit-assignment challenge which delays rule learning.

## Results

### Monkeys are slower rule learners than humans

We compared the ability of monkeys and humans to rapidly adapt to changes in task contingencies and learn new rules in a modified version of the WCST. On each trial, subjects were presented with 4 objects and received feedback upon selecting one of them (Fig. 1a, middle). Each object was composed of one unique feature from each of three dimensions - visual pattern, shape and color (Fig. 1a, top). Every feature appeared in one of the objects on each trial, but object compositions changed across trials. On a given block of trials, subjects received positive feedback (monkeys: food reward; humans: the word “CORRECT” displayed on screen) for selecting the object with one of the twelve features (e.g. red) — the current hidden rule — and negative feedback (monkeys: timeout; humans: the word “INCORRECT” displayed on screen) otherwise. After subjects demonstrated that they had learned the current rule by reaching criterion performance on the current block, a new block was initiated through an uncued switch to a randomly chosen new rule (Fig. 1a, bottom). Parameter differences in the task implementation for the two species, including the response type, trial epoch durations and learning criteria, are outlined in Supplementary tables 1 and 2.

Remarkably, well-trained monkeys learned new rules within only tens of trials. Yet, they were over three times slower than humans (Fig. 1b; monkeys: 27.84 *±* 2.92 trials (mean *±* SEM); humans: 5.98 *±* 0.52 trials), a learning deficit that was significant following a correction for the inter-species difference in the learning criterion (Supplementary Fig. 1; monkeys: 20.61 *±* 1.52 trials; humans: 5.98 *±* 0.52 trials; bootstrap test with *t*-statistic, p *<* 0.005). We then sought to explain the inter-species computational differences that produce this rule learning slowing in monkeys. Specifically, we focused on inferring individuals’ rule-learning strategies from behavior, and identifying the species differences that contribute to the learning speed difference. One possible strategy, Win-Stay Lose-Shift (WSLS), is widely-reported during rule learning in many species, particularly in the two-armed bandit problem where the identity of the more rewarding arm must be learned and can change over trials. Here, one of the arms is repeatedly chosen as long as this produces positive feedback (win-stay). When negative feedback is received the other arm is chosen on the next trial (lose-shift). This strategy can be cast as a decision process comprised of two behavioral states — persist and avoid — where the choice of the currently rewarded arm is in the persist state and it transitions to the persist or avoid states subject to positive or negative feedback, respectively, while the choice of the other arm is in the avoid state and transitions to the persist state when that arm was not chosen on the previous trial and negative feedback was received (Fig. 1c).

The WSLS strategy is computationally efficient, and requires that the subject attend to and maintain only a single arm’s identity in working memory. By replacing arm identity with feature identity, the approach is readily adapted to solve WCST problems and always finds the rule. However, feedback is equally informative about all three features in the chosen object, not just the attended one. Due to this neglect of information about unattended features, a simulated WSLS agent learns rules much slower than optimal: indeed humans learn more rapidly (Supplementary Fig. 1; WSLS agent mean: 13.31 trials, std. dev.: 12.85 trials; humans: 5.98 *±* 0.52 trials). This underscores a tradeoff between computational and information efficiency in multi-dimensional environments. Simultaneously maintaining and updating beliefs about multiple features is more information efficient, but increases computational complexity and working memory demands. In contrast, attending to a single feature at a time is computationally simpler but inefficient in its integration of trial outcomes. In the following sections, we address how the two species solve this tradeoff.

### Dynamic model uncovers hidden states during rule learning

Prior cognitive model comparisons of human behavior in WCST variants provide evidence for rule-learning strategies wherein subjects selectively attend to and learn about individual features or dimensions, rather than feature configurations (i.e. objects) [17, 18, 20]. It is argued that such a mental representation of stimuli in terms of features resolves the curse-of-dimensionality which impairs learning efficiency in high-dimensional environments. For example, it is far more efficient to learn the value of 12 features than the dozens of objects they can be combined into. Drawing on these findings, we developed a behavioral model to predict the probability of choosing individual features given their choices and outcomes on previous trials. However, in contrast to earlier work, our model does not postulate a specific internal belief structure and update rule, thus making fewer assumptions regarding the learning algorithm underlying a subject’s behavior. Instead, it aims to discover in an unbiased manner how the decision making process evolves as a function of feedback. Recently, this approach has been successful at revealing previously unobserved behavioral states underlying human, rodent and fruit-fly decision making [22, 23, 24].

For each feature, we model whether the feature is chosen or not (denoted as *c*) as a function of past choices and their outcomes (*h*) via a Bernoulli Generalized Linear Model (GLM) (Fig. 2a; see *Methods*). The choice outcome on an earlier trial is represented by a one-hot four dimensional vector where the dimensions represent whether positive feedback was received after choosing the feature on the trial (*C*^+^), negative feedback was received after choosing the feature (*C*^−^), positive feedback was received after not choosing the feature on the trial (*NC*^+^), or negative feedback was received after not choosing the feature (*NC*^−^). This allows us to assess separately how the present choice depends on past choice outcomes both when that feature was chosen (direct) and when it was not (indirect). Furthermore, the model permits dynamic changes in how past choices and outcomes are transformed into a present choice via hidden states (*s*). A feature’s associated hidden-state also undergoes a transition at the end of each trial depending on past choices and outcomes, which may reflect updates to the feature’s value based on past choice outcomes, or a change in the level of attention to the feature, or even a shift in strategy (i.e. how a feature’s history is weighted in determining its choice). Note that while the model permits these possibilities and others, it does not prescribe the nature and function of the states. Rather, the states and their dynamics emerge upon fitting the model to behavioral data. These hidden state dynamics are modeled as an input-dependent or Input-Output Hidden Markov Model (IOHMM) [25].

**Figure 2:**
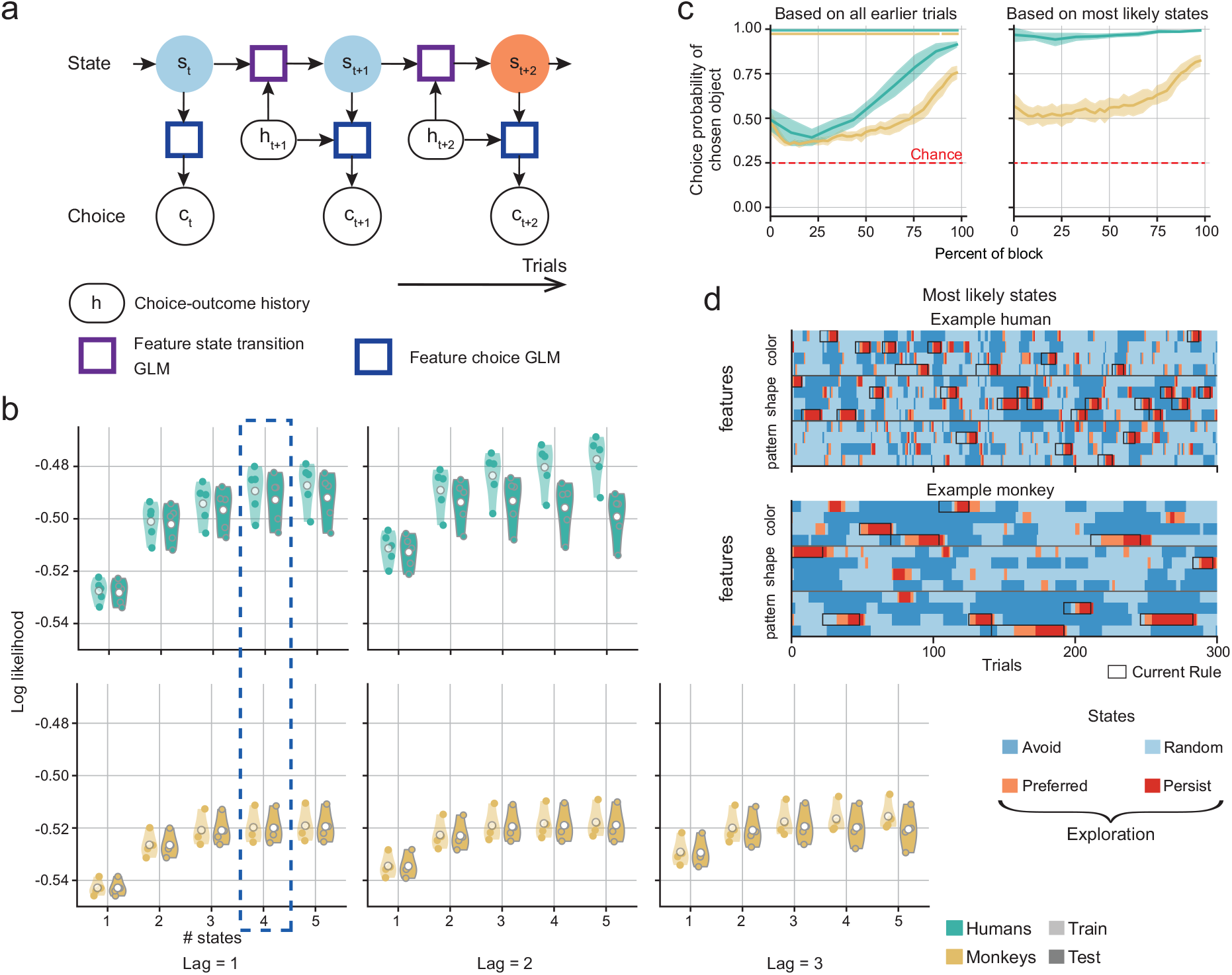
IOHMM-GLM model fits uncover dynamic changes in choice behavior during rule learning. **a**. IOHMM-GLM model architecture fit to data. The model predicts the choice of a feature c at each trial t from the choice-outcome trial history h via a GLM. Hidden states s determine the GLM’s parameters. These states can transition at each trial also based on the choice outcome trial history via a separate state transition GLM. **b**. Model fit log-likelihoods on training and test datasets for each human (green) and monkey (brown) subject in models with varying numbers of states and that use choice outcomes from varying numbers of previous trials (lag) to determine the feature choice and state-transition probabilities. Each point represents a single subject’s mean over a 5-fold cross-validation and over 5 (monkey) or 10 (human) different model initializations. Each subject’s best-fit model with 4 states and lag 1 (dashed blued box) was chosen for further analysis. **c**. Probability of selecting the chosen object produced by a model extension based on feature choice probabilities predicted only from choice outcomes on earlier trials (left), and on feature choice probabilities computed from most-likely state estimates derived from past, present and future choice outcomes (right). The probability on each trial was binned according to the trial’s relative position in the rule block and averaged across blocks. Line and shading represent the mean and standard deviation across subjects for each species. Dots represent block percentiles at which the average object selection probability is significantly above chance (bootstrap test with t-statistic, p < 0.05). **d**. Most-likely states estimated by the model for 300 trials in an example human (top) and monkey (bottom) subject. The rule on each block is outlined in black.

The IOHMM-GLM’s goodness of fit to behavior depends both on the number of previous trials determining a subject’s choice (lag), and on the number of possible hidden states. Accordingly, we fit IOHMM-GLMs to each subject’s behavior while systematically varying these two parameters (Fig. 2b). Across subjects in both species, model accuracy showed a stronger dependence on the number of states than lag. Crucially, accuracy plateaued as the number of states increased, and exhibited over-fitting at higher lags. In this task, subjects choose objects rather than individual features. Therefore, we extended our model to compute the probability of choosing each object in a trial, based on the model’s predicted probability of choosing individual features on that trial (see *Methods*). Fits of this model extension to each subject’s behavior based on each of the IOHMM-GLMs in Figure 2b revealed a qualitatively similar relationship between model accuracy and the underlying parameters (Supplementary Fig. 2a). For each subject, the best-fit model comprised of 4 states and lag 1 (history from the previous trial only) does not overfit the data while producing prediction accuracies at or very close to the performance plateau. Therefore, we selected these models (dashed-blue box, Fig. 2b, Supplementary Fig. 2a) for further analysis.

Figure 2c (left) shows the choice probability predicted by these 4 state lag 1 models for the chosen object at each trial after a rule switch, averaged over rule blocks; averaging across blocks is achieved by normalizing the trial number by the block length. The results show that the model’s prediction of the chosen object is significantly above chance (0.25) in both species (monkeys: 0.47 *±* 0.02; humans: 0.63 *±* 0.02). Also, prediction accuracy improves as the rule is learned over the block’s time course. Our primary goal, however, is to find the most accurate explanation for each subject’s rule learning behavior, rather than predict their future choices. For this, we consider the most likely sequence of states across trials inferred by the model for each feature. Rather than predict the most likely state on each upcoming trial given only past choice outcomes, the most likely sequence of states is the maximum a posteriori probability (MAP) estimate of the sequence of states across all trials in an experimental session — each estimated or inferred state best explains not only the present choice but also past and future choices subject to the model’s choice probabilities in the inferred past/future states and its state transition probabilities between the inferred present and past/future states (see *Methods*; Figure 2d). The model is generally quite confident in its MAP estimates of the most likely sequence of states, as evidenced by the the cumulative density of their posterior probabilities (Supplementary Fig. 2b). Moreover, since the inferred states for each feature are estimated from past, present and future choices, they yield more accurate estimates of the choice probabilities for chosen objects (Fig. 2c, right). For this reason, we rely on the inferred states to identify the rule learning strategy in each species and to interpret the inter-species differences therein.

To gain insight into the interpretation of our model fits, we similarly analyzed the choices of a simulated Win-Stay Lose-Shift agent (Supplementary Fig. 1). By construction, we know that the model’s choices only rely on the previous trial. As expected, higher lag models tend to overfit the agent’s choices (Supplementary Fig. 3a). While the agent’s true behavior has only two states (Fig. 1c), we find that a three-state model provides a better fit. Our model splits the avoid state into two states — random and avoid. This is due to a combination of the task’s structure and our model setup. By picking one feature consistently across trials, the agent necessarily avoids other features in the same dimension. However, the agent’s choices of features in other dimensions appear random, since an object is composed of one feature from each dimension and objects compositions are generated randomly on each trial. Thus, the appearance of this additional state results from our model’s treatment of each feature independent of its relationship to other intra-dimensional features, a simplifying assumption that allows for tractable fitting. Nevertheless, the model largely recovers the hidden states and state-transitions that drive the agent’s behavior — it correctly identifies when a feature is associated with the persist state 57.6% of the time and accurately determines the underlying decision process (Supplementary Fig. 3c-d). Collectively, these results show the reliability of this modeling approach to explain rule-learning in both species.

### Hidden states reflect feature-based attention and reveal qualitatively similar strategies in the two species

Learning is often conceptualized as updates to a decision-making schema based on past decisions and their outcomes [18, 26]. We sought to identify hidden states that capture this decision-making process and to explore what they reveal about the *dynamics* of human and monkey rule learning in the WCST. In both humans and monkeys, a comparison of the choice probability of features associated with each state — calculated by marginalizing the model’s predicted choice probability under each state and history (Supplementary Fig. 4a) over the choice outcome histories (Supplementary Fig. 4b) — revealed that the model determines states based on distinct probabilities of choosing the associated feature, ranging from below chance (*avoid*) to chance (*random*) to above chance (*preferred*) to very high (*persist*) (Fig. 3a). That is, the model states correspond to levels of attention paid to each feature. Moreover, this result was consistently observed in the majority of the models fit to the behavior in both species, as well as in a simulated WSLS agent (Supplementary Fig. 3). Since features associated with the preferred or persist state are favored during rule-learning, we refer to them as being under exploration. We will show that the estimation of the attentional state towards each feature at each trial permits a systematic analysis of when features are selected for or withdrawn from exploration, and how the choice outcome history informs these decisions. This exercise fosters an exposition of the decision-making process that describes the rule learning strategy in both species, the resulting learning dynamics between rule switch and rule learning, and the identification of those differences in the decision-making process that most prominently explain the learning performance difference between the two species.

**Figure 3:**
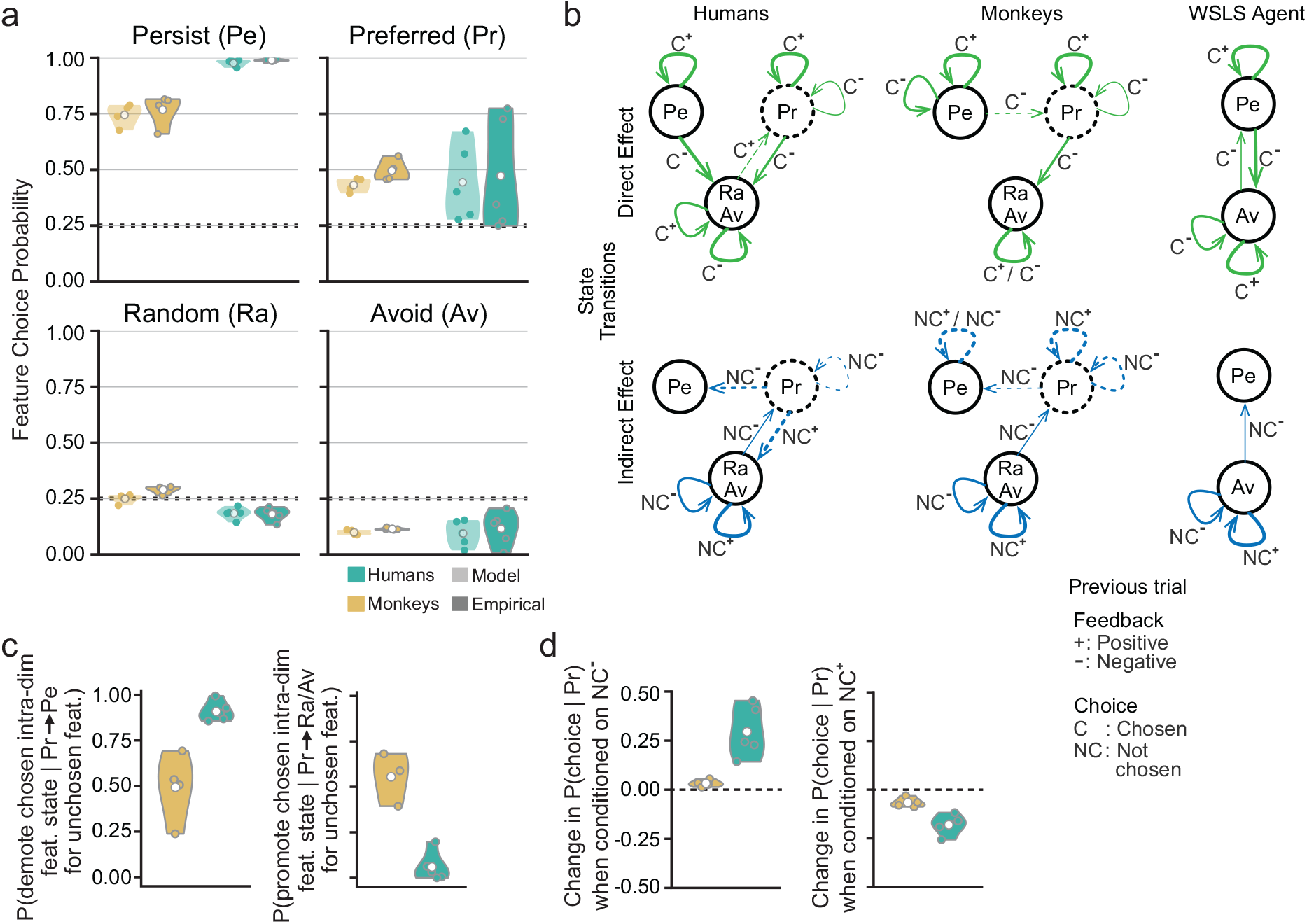
Model describes rule-learning dynamics in terms of changes in feature-attentional states. **a**. Choice probability of features associated with each state in human (green) and monkey (brown) subjects computed directly from model parameters and measured empirically based on most-likely state estimates. Choice probabilities order feature states akin to levels of attention. **b**. Decision process describing how humans, monkeys and the WSLS agent start, continue and stop exploring a feature, derived from their history-dependent state transition probabilities. Process is decomposed based on outcome-dependent transitions when the feature is chosen (direct effect) or not chosen (indirect effect). Arrow thickness indicates probability of the transition. Dashed lines highlight deviations from the WSLS strategy. **c**. Probabilities of demoting (left; promoting, right) the state of a chosen feature to a state with higher (lower) choice probability when an unchosen intra-dimensional feature is promoted (demoted) from the preferred to persist (random/avoid) state. Measurements test to what extent indirect effects of promoting or demoting features in the preferred state result from changing the state, and therefore the choice probability, of a chosen intra-dimensional feature. Perfect causality would coincide with a probability of 1.0. **d**. Change in the choice probability of a feature in the preferred state after after receiving negative (left; positive, right) feedback for choosing a different feature. The indirect effect significantly increases (decreases) the feature choice probability.

Since these analyses rely heavily on the most-likely state estimates, we validated the consistency of these estimates with the model’s parameters. First, we compared the feature choice probability per history and state computed directly from the fit parameters (model) and measured based on the state estimate for each feature on each trial (empirical). The two measurements yield consistent results demonstrating that the most-likely states are estimated not only to best explain the sequence of choices but also to conform with the model’s parameters. Next, we similarly compared state transition probabilities per history computed directly from the fit parameters (model) and measured based on the state estimate for each feature on each trial (empirical)(Supplementary Fig. 5). Here again, we find that the transition probabilities computed from the fit parameters (model) are consistent with empirical measurements of the transition statistics based on the most-likely state estimates. Figure 3b schematizes the decision process in the two species derived from their state transition probabilities. The thickness of an arrow indicates the probability of the respective transition; extremely rare transitions have been pruned. Similar to the WSLS agent, we find that a feature is most often associated with the avoid state while an intra-dimensional feature is simultaneously under exploration (Supplementary Fig. 4c). Since the avoid state likely emerges due to this interdependence between the choices of inter-dimensional features, which our model forgoes for tractability, we do not treat it as distinct from the random state.

We would like to compare observed behavior in the WCST with a WSLS strategy. To do so, we must take into account task structure differences between the WCST and the 2-armed bandit task (Fig. 1c).The composition of a chosen object by three features forces the choice of features in the avoid state of the WSLS strategy. Thus a WSLS decision process for the WCST must define transitions for such features when they are chosen. Moreover, the multi-dimensional environment of the WCST offers multiple alternatives for a subject to shift attention to during lose-shift, compared to the 2-armed bandit task. An updated WSLS strategy that accounts for these differences is depicted in Figure 3b (right). We can now compare the decision process inferred by the model for the two species (Fig. 3b, left-middle) to this WSLS decision process, revealing salient differences that are delineated by dashed lines. Key among these is the existence of a *preferred* state where items are not chosen with certainty (or near-certainty) as in the *persist* state, but above chance. The effect of direct feedback (as a result of choosing the feature) on these states and the random/avoid states is similar to those in the WSLS decision process. For example, both species select a feature in the random/avoid state for exploration (by promoting it to the preferred state) seemingly at random after receiving negative feedback for choosing other features. However, an interesting exception is that humans sometimes choose to explore such a feature upon receiving positive feedback for choosing it.

Larger differences emerge with regards to indirect effects of feedback. A feature may not be chosen on a trial when it is associated with the preferred state (feature choice probability in preferred state *<* 1, Fig. 3a). However, its state may still transition subject to the feedback received at the end of the trial — an indirect effect. For example, humans and, to a lesser extent, monkeys demote features from the preferred to the random/avoid state upon receiving positive feedback for choosing a different feature. Consequently, their probability of subsequently choosing an unchosen feature that was associated with the preferred state decreases (Fig. 3d, right). They also promote features from the preferred to the persist state upon receiving negative feedback for choosing a different feature. Consequently, their probability of subsequently choosing an unchosen feature that was associated with the preferred state increases (Fig. 3d, left). This is striking because it is the only way a feature can transition into the persist state, which appears to be reserved mainly for a feature that the subject determines to be the rule (Fig. 2d). Receiving positive feedback for choosing a feature in the preferred state does not definitively confirm that it is the rule, since the rule may be among the other two features in the chosen object. Confidence in the rule’s identity may be increased based on the consistency of receiving such direct positive feedback across many trials. Alternatively, it may be done by ruling out other candidates, that is, after receiving negative feedback for choosing an object with a different candidate feature. Consistent with this interpretation, measurements show that when a feature under exploration is not chosen, the object that is chosen often contains a different feature that is also under exploration (Fig. 4d).

**Figure 4:**
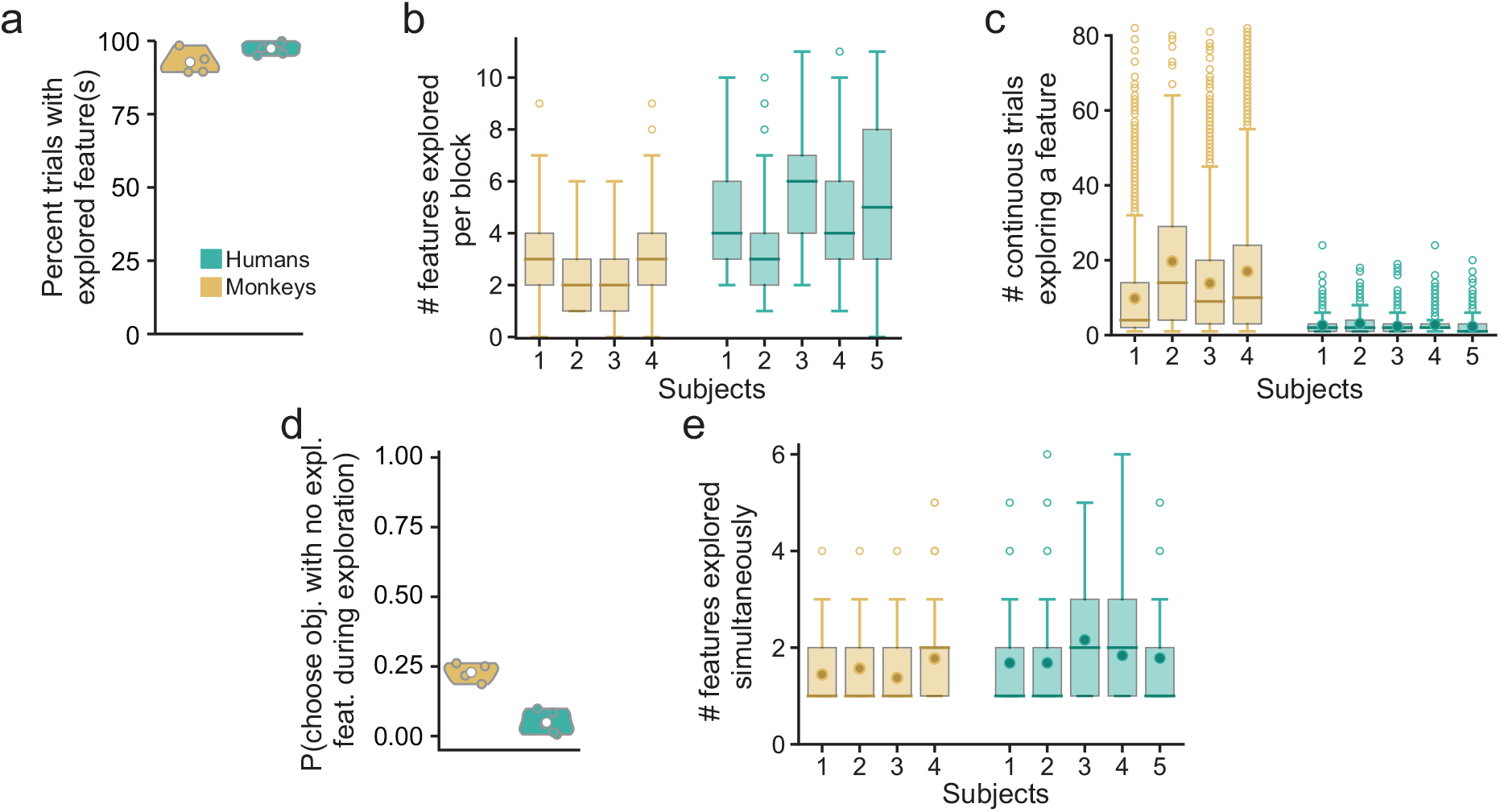
Monkeys and humans explore multiple features for several trials in a row to evaluate them. **a**. Percent of all trials where at least one feature is under exploration by humans (green) and monkeys (brown). **b**. Distribution of the number of features explored by each monkey and human subject in a block. **c**. Distribution of number of continuous trials with a feature in an exploration state. **d**. Probability of choosing an object with all features in the random or avoid state, while at least one other feature is in the preferred or persist state. **e**. Distribution of number of features simultaneously explored by each monkey and human subject in trials where at least one feature is under exploration.

This approach of promoting a feature to the persist state as an inferred consequence of ruling out an alternative candidate, rather than integrate direct positive feedback across trials in favor of the feature, may be favored by both species due to its computational simplicity — it relies on the outcome of just the previous trial rather than multiple trials and thereby reduces working memory demands. It is possible that these inference-like computations are not deliberate but an inadvertent consequence of demoting or promoting an intra-dimensional chosen feature. For example, given that the probability of choosing all shapes must sum to 1, when one shape is demoted after its choice produces negative feedback, the probability of choosing another shape that was associated with the preferred state may automatically increase, forcibly promoting it to the persist state. Measurements of the probability of demoting (promoting) the chosen feature while promoting (demoting) an unchosen intra-dimensional feature in the preferred state are mixed: monkeys do so at chance levels; humans always (seldom) demote (promote) the chosen feature (Fig. 3c). Nevertheless, these indirect-effect transitions directly and significantly alter the subsequent choice probability of the unchosen feature (Fig. 3d). In summary, the best-fit models discover feature-based attentional states whose dynamics show marked deviations from a Win-Stay Lose-Shift strategy.

### Both species simultaneously evaluate multiple features over several trials during rule-learning

The explore-exploit dilemma pits the benefit of continuing to select a recently rewarded option (exploit) against the benefit of selecting a different and potentially more rewarding (but possibly less rewarding) option (explore). While much work has been done to determine how humans and other animals navigate this dilemma [27, 28, 29], how they deal with it in a multi-dimensional environment with transiently overlapping options remains unclear. Which of the three features of a chosen and rewarded object should be exploited on the next trial, given that they are unlikely to appear co-located in the same object on the following trial? How should the tradeoff between the computational complexity and information efficiency of exploring several features at once be resolved?

The model finds that both species continuously explore one or more features (Fig. 4a). In the process, they explore multiple features over the course of a block before ultimately identifying the rule (Fig. 4b). Moreover, each feature is often explored for a series of several trials in both species (Fig. 4c). But the number of these trials is substantially larger in monkeys, a finding we analyze more closely in the following sections. The model also indicates that both species often explore multiple, but not all, features at a time (Fig. 4e). This is consistent with the theory of selective attention [30, 31] wherein objects are selectively attended to (or filtered for higher processing) subject to an internally-maintained set of relevant perceptual features (or attentional filters). It also underscores the solution of both species to the computational complexity-information efficiency tradeoff. Since it is computationally challenging to simultaneously attend to and evaluate all twelve features over several trials but inefficient to attend to one feature at a time, both species evaluate a small subset of all features at a time. Indeed, during exploration it is uncommon for either species to select an object where none of the features is in the preferred or persist states (Fig. 4d). However, monkeys do engage in such random exploration much more frequently than humans.

From these results, we conclude that both species exhibit a deliberate form of exploration to address the challenges inherent in the task environment. Features, often more than one at a time, are selected for exploration via promotion to the preferred state after choices of other features produce negative feedback (Fig. 3b). They are then continuously explored so long as they produce positive feedback, until alternatives are ruled out at which point they are then promoted to the persist state, or they are themselves ruled out after choosing them produces negative feedback (or, in the case of humans, choosing other features produces positive feedback).

### Categorization of feature attentional states characterizes learning dynamics

Rule learning proceeds through a sequential process that progressively reduces ambiguity regarding the rule’s identity until it is ultimately determined. Our model reveals elementary feature-specific computations that individuals in each species apply to maintain and update a small subset of candidate features, one of which may determine the rule. In order to elucidate the resulting over-arching learning dynamics that governs the sequential rule learning process, we developed a simple approach to categorize individual trials based on the features under exploration and the true rule (Fig. 5a). The categories are mutually exclusive and exhaustive – each trial falls into one and only one category. These are:

**Figure 5:**
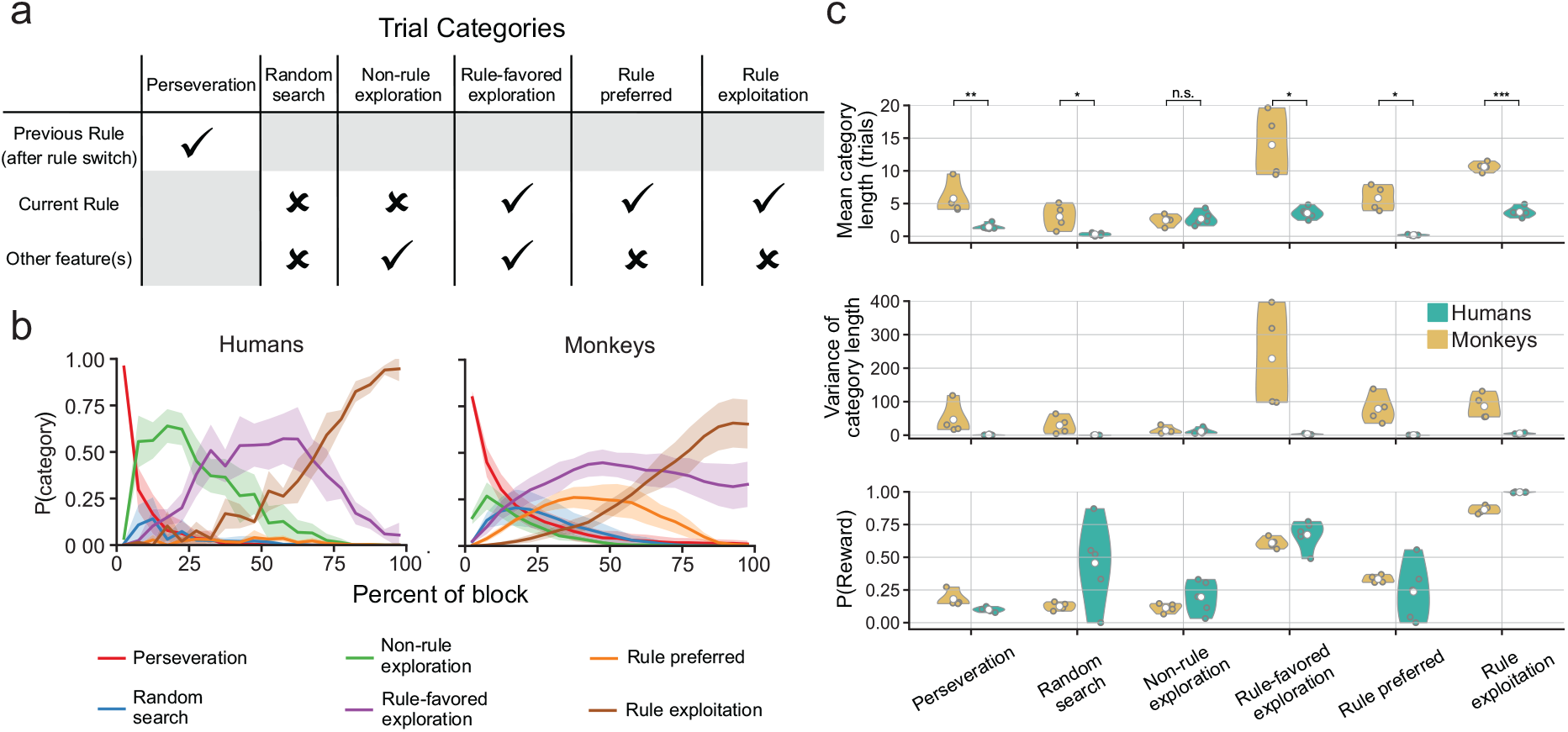
Exploration-based trial categories reveal learning dynamics and identify causes for monkey learning performance deficit. **a**. Definition of the six trial categories based on whether the features under exploration during the trial include the rule feature. **b**. Distribution of trial categories at each percentile of rule block. Lines (shaded areas) reflect mean values (standard errors of the mean) across subjects. **c**. Trial category summary statistics (top: mean number of trials; middle: variance of number of trials; bottom: reward probability) across rule blocks for human (green) and monkey (brown) subjects. Inter-species comparisons of the mean number of trials per category reveal significant differences in the perseveration, random search, rule-favored exploration, rule preferred and rule exploitation categories (bootstrap test with t-statistic); n.s. - not significant; * p < 0.1; ** p < 0.01; *** p < 0.001.

- “perseveration”: a continuous series of trials following a rule switch when the feature governing the previous rule is associated with the persist state,
- “random search”: trials when none of the features are under exploration (i.e. associated with the preferred or persist states),
- “non-rule exploration”: trials when one or more features are under exploration not including the rule feature,
- ““rule-favored exploration”: trials when one or more features including the rule are under exploration,
- “rule preferred”: trials when only the rule is associated with the preferred state,
- “rule exploitation”: trials when only the rule is associated with the persist state.

We compare the distribution of categories for trials across the course of a rule block between species (Fig. 5b). Humans show a progression from perseveration to non-rule exploration, where non-rule features are explored and ruled out, to rule-favored exploration, where the rule feature is simultaneously explored with non-rule features, to rule exploitation, once other candidates are ruled out and the rule is identified. Monkey rule learning is described by similar dynamics except for a much higher incidence of the rule preferred category for most of the block and a larger (smaller) proportion of the blocks ending in the rule-favored exploration (rule exploitation) category. That is, a significant subset of blocks end with the model indicating that the monkey is simultaneously exploring the rule and at least one other non-rule feature, even if the monkey has met a learning criterion. These results show that our categorization approach expresses human and monkey rule learning dynamics in terms of behaviorally interpretable learning stages; for example, an increase in the reward rate following a rule switch in both species is marked by the onset of rule exploration with the rule-favored exploration category (Fig. 5c, bottom).

Examining the number of trials spent in each category determined bottleneck categories that produce the rule learning performance deficit in monkeys (Fig. 5c). Specifically, monkeys spend much longer perseverating on the previous rule, in disambiguating the rule feature from non-rule features (rule-favored exploration) and demonstrating that they have learned the rule (rule preferred or exploitation). The latter two sources of the learning performance deficit in monkeys also explain a majority of the variance in their performance across blocks (Supplementary Fig. 7, bottom). In contrast, the number of trials humans spend exploring non-rule features before selecting the rule feature for exploration (non-rule exploration) largely determines the variance in their rule-learning performance.

### Random exploration prolongs the expression of learning in monkeys

A key difference between the two species identified via this trial categorization is that monkeys spend many more trials than humans in the rule preferred or exploitation categories. These extra trials spent demonstrating or expressing that the rule has been learned significantly increases both the block length mean and variance (Fig. 5c, Supplementary Fig. 7). The inter-species difference in learning criteria explains a portion of the difference in the mean length of the rule exploitation category. However, the remaining difference in the mean length as well as the difference in the variance of the category’s length is unexplained. We hypothesized that the larger mean and variance of the rule exploitation category’s length in monkeys compared to humans (Fig. 6a) may result from their random exploration of other features when a feature is already associated with the persist state (Fig. 3a). This behavior is unique to monkeys and is prevalent even during rule exploitation trials (Fig. 6b) — after they have identified the rule, monkeys occasionally choose objects that do not include the rule feature.

**Figure 6:**
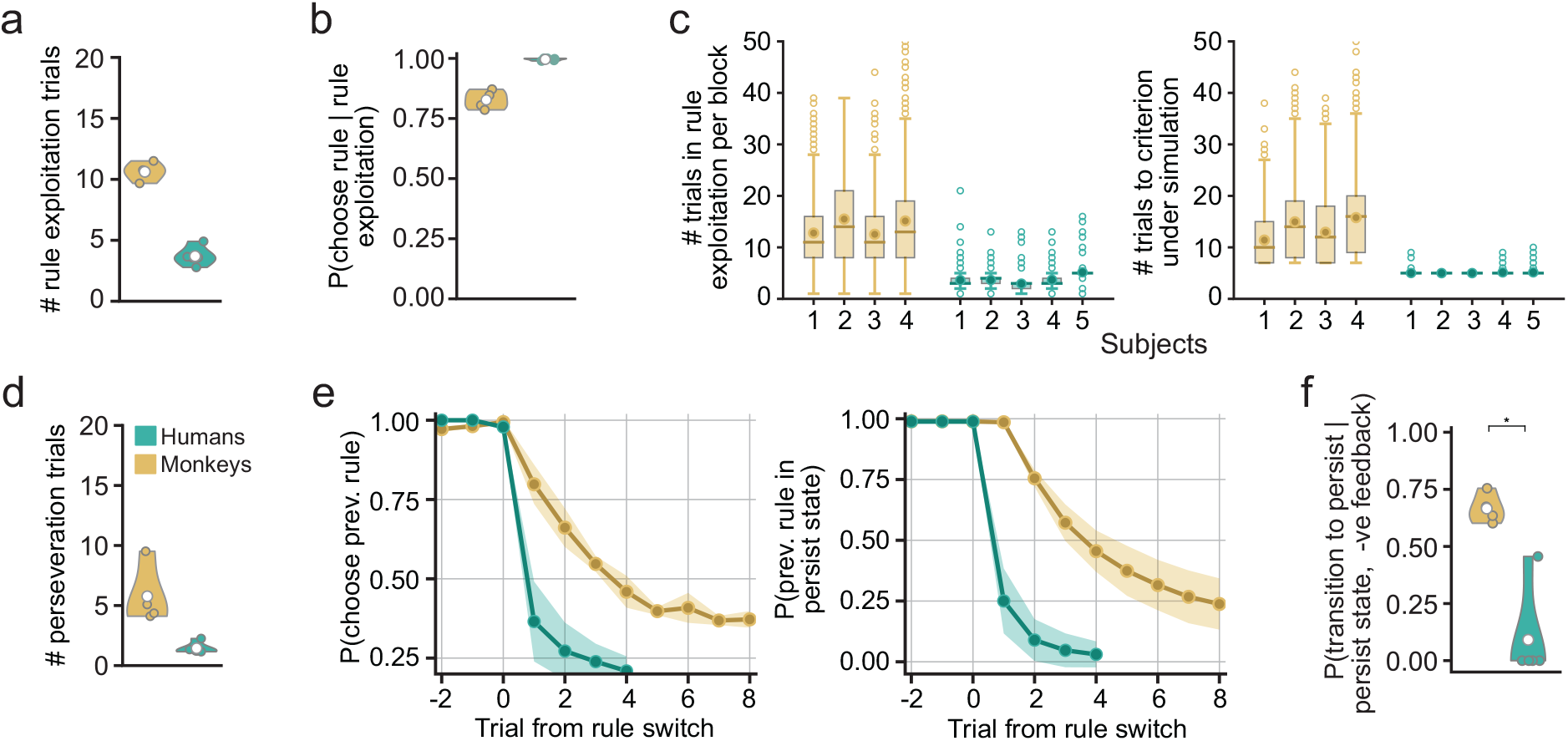
Random exploration and perseverative errors prolong monkey rule learning. **a**. Mean number of trials spent by human (green) and monkey (brown) subjects in the rule exploitation category per rule block. **b**. Probability of selecting an object with the rule feature across trials in the rule exploitation category. Monkeys occasionally explore other objects compared to humans. **c**. Distribution of the number of trials spent by human and monkey subjects in the rule exploitation category per rule block (left), and by simulated agents that select the rule feature with probabilities in (b) until they reach a learning criterion (right). **d**. Mean number of trials spent by human and monkey subjects in the perseveration category per rule block. **e**. The probability of humans and monkeys choosing the previous rule feature at each trial after a rule switch (left) is commensurate with the probability of the previous rule feature being associated with the persist state (right). **f**. The probability of the previous rule feature transitioning back into the persist state after its selection produces negative feedback is higher in monkeys (bootstrap test with t-statistic); * p < 0.1.

To test our hypothesis, we simulated agents that select the rule feature with the same probability as monkeys and humans do during the rule exploitation category and asked how many trials it would take these agents to reach a learning criterion. The results revealed that the trial count distributions of the simulated agents were nearly identical to the corresponding subjects (Fig. 6c), thus confirming our hypothesis. Similar “random errors” have been observed in humans with focal lateral prefrontal lesions on the WCST [19], where they were attributed to distraction, or a failure to maintain the rule in working memory. However, it remains unclear whether the monkeys in our experiments were more distractable than their healthy human counterparts, or deliberately adopted occasional random exploration as part of their strategy — for example to prolong a highly rewarded state.

### Reduced sensitivity to negative feedback increases perseverative errors in monkeys

Perseverative errors occur when the feature that governed the rule on the previous block continues to be chosen following the rule switch despite receiving negative feedback for the choice. These errors are characteristic of frontal lobe damage and dysfunction [10, 21] and are believed to reflect a cognitive deficit in adapting to changes in task contingencies. Pronounced perseveration error rates are also observed in patients with neuropsychiatric [32, 33, 34] and substance abuse disorders [20, 33]. Interestingly, our model’s association of the persist state with the previous rule feature during consecutive trials immediately following a rule switch suggests that monkeys persevere on the previous rule for several more trials than humans (Fig. 6d). Indeed, direct measurements showed that the probability of of choosing the previous rule after a rule switch is consistent with such a state estimate in both species (Fig. 6e).

In order to determine the cause underlying the elevated perseveration in monkeys, we asked which choice outcome(s) most explained the difference in continued persistence with the previous rule between the two species. The analysis showed that humans were far more likely to demote the chosen previous rule feature from the persist state in response to negative feedback compared to monkeys (Fig. 6f). Monkeys’ weaker sensitivity to negative feedback parallels that of humans with substance abuse disorders and prefrontal lesions, who also persevere more than healthy controls [20, 21].

### Reduced negative feedback sensitivity compromises efficient credit assignment and prolongs rule learning in monkeys

The largest contribution to the inter-species difference in rule learning performance is from trials in the rule-favored exploration category where the rule feature is concurrently explored with one or more non-rule features (Fig. 7a - Fig. 7b, left). While it is reasonable to explore the rule for several consecutive trials as it produces rewards, what must be explained is why non-rule features are concurrently explored for many more trials by monkeys. Indeed, monkeys continuously explore individual non-rule features for many more trials during the rule-favored exploration category (Fig. 7b, right). This explains the lengthier duration of the category in monkeys, and is caused by a higher probability of a non-rule feature transitioning back into an exploration state during rule-favored exploration trials (Fig. 7c). Analysis further showed that this inter-species difference in transition probability is explained by a lower sensitivity of monkeys to either form of negative feedback – direct, when the non-rule feature is chosen and negative feedback is received, and indirect, when it is not chosen and positive feedback is received (Fig. 7d).

**Figure 7:**
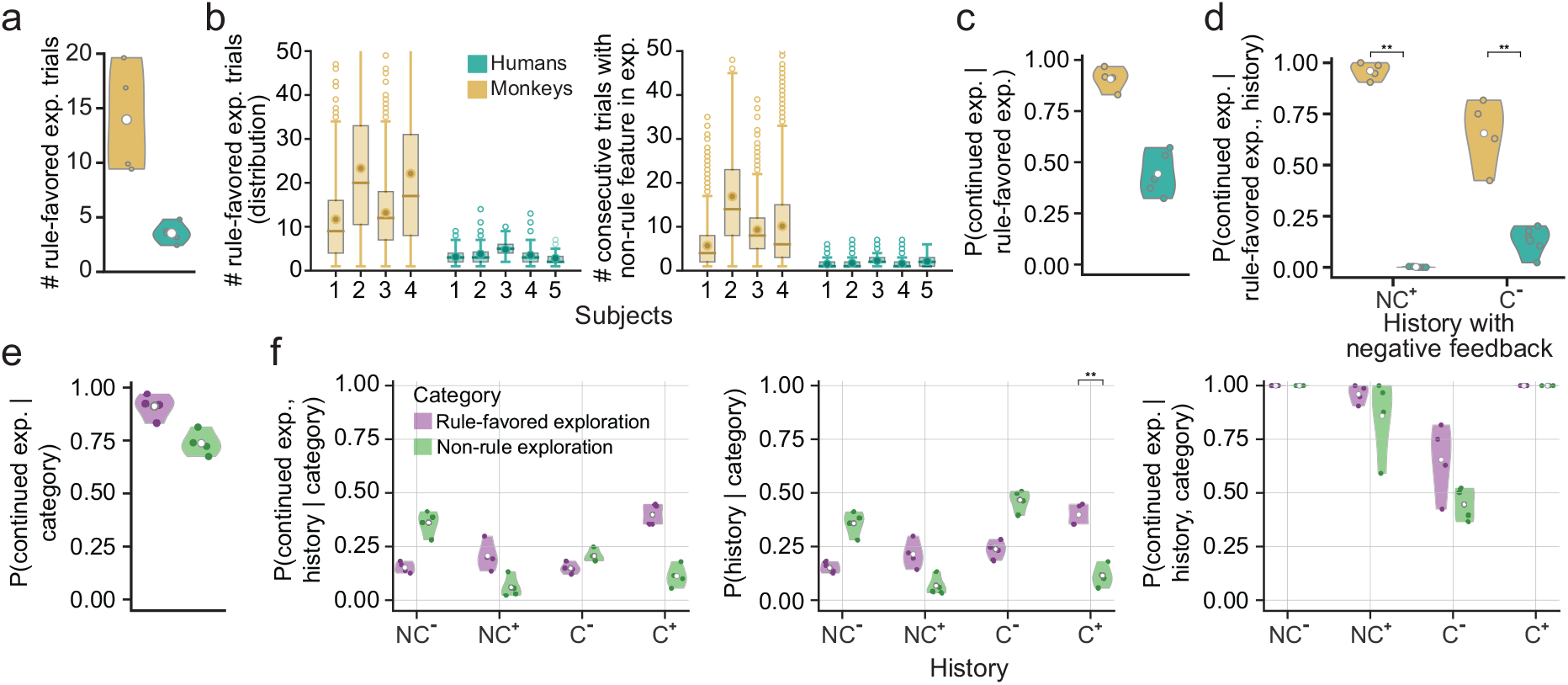
Diminished negative feedback sensitivity prolongs concurrent exploration of rule and non-rule features. **a**. Mean number of trials spent by human (green) and monkey (brown) subjects in the rule-favored exploration category per rule block. **b**. Distribution across rule blocks of the number of trials spent in the rule-favored exploration category by each subject (left), and of the number of consecutive trials spent by them exploring individual non-rule features during this category (right). **c**. Probability of a non-rule feature transitioning back into an exploration state during rule-favored exploration trials. **d**. The probability of a non-rule feature transitioning back into an exploration state upon receiving negative feedback for choosing it (direct negative feedback) or positive feedback for choosing a different feature (indirect negative feedback) during rule-favored exploration trials is higher in monkeys (bootstrap test with t-statistic). **e**. Probability of a non-rule feature transitioning back into an exploration state during rule-favored exploration trials and non-rule exploration trials in monkeys. **f**. Joint probability of a non-rule feature transitioning back into an exploration state and each choice outcome history occurring during rule-favored exploration trials and non-rule exploration trials in monkeys (left); Probability of each choice outcome history occurring during either category (middle); Probability of a non-rule feature transitioning back into an exploration state in response to each choice outcome history during either category (right). The higher probability of a non-rule feature transitioning back into an exploration state during rule-favored exploration trials compared to non-rule exploration trials is explained by a higher incidence of direct positive feedback for choosing the non-rule feature in the former category (bootstrap test with t-statistic); ** p < 0.01.

While both species also explore non-rule features during non-rule exploration trials, this category is relatively short in both humans and monkeys (Fig. 5c). So what explains the difference in duration between the two categories in monkeys? Since the non-rule exploration category is followed by rule-favored exploration trials, one possibility is that it is cut short by the onset of exploration of the rule feature as the non-rule feature continues to be concurrently explored for many more trials. However, measurements in monkeys showed that non-rule feature exploration only occasionally spans the two categories (probability = 0.27% *±* 0.04%). Therefore, the number of trials during which a non-rule feature is explored by monkeys is usually much smaller when it happens in the non-rule exploration category than in the rule-favored exploration category.

This difference in duration is reflected in a higher probability of a non-rule feature transitioning back into an exploration state during rule-favored exploration trials (Fig. 7e). Since the probability of transitioning back into the explore state is a marginalization of its joint probability with the choice outcome it follows, we asked what choice outcome history best explains the transition probability difference between the two categories. Measurements showed that receiving positive feedback for choosing the non-rule feature (*C*^+^) is the key differentiator between the joint probabilities for the two categories (Fig. 7f, left). This could either be because the transition probability in response to this history is different under the two categories, or because the frequency of the history is different under them. We found that while the transition probabilities are similar (Fig. 7f, left), monkeys are more likely to have received positive feedback for choosing a non-rule feature under exploration during the rule favored, exploration category (Fig. 7f, middle). When the rule feature is concurrently explored with a non-rule feature (rule favored, exploration) the probability of selecting them both when they co-locate in an object is higher. This increases the probability of receiving positive feedback for choosing the non-rule feature, which makes appropriately assigning credit to the rule feature challenging. This underscores the importance of negative feedback sensitivity in demoting non-rule features from exploration states, in the absence of which the duration of concurrent exploration of the rule and non-rule feature(s) is prolonged.

## Discussion

Methodological and technological advances in training and recording from animal models now allow for the study of increasingly complex behaviors in non-humans. However, before interpreting their brain activity as a human-like model of neural computation, it is important to ascertain whether their computational algorithms are human-like. Indeed macaque monkeys and humans learn the structure of tasks in different ways (monkeys via impoverished reward-based feedback, and humans via rich verbal instruction plus feedback), raising the possibility that while they both learn the same tasks, they may enlist different abstractions, cognitive operations and neural mechanisms [2]. Our study aims to assess whether macaques and humans employ similar mental representations and operations to perform a cognitively complex task that relies on several interdependent cognitive processes, such as the Wisconsin Card Sorting Test. The results of such studies can play a crucial role in interpreting inter-species comparisons of the neural correlates of these representations and operations. Our findings demonstrate that both species employ similar overall strategies to perform the task (Fig. 3a-b). However, key differences in the decision criteria of these strategies explain monkey performance deficits on the task.

The Wisconsin Card Sorting Test was originally developed to test cognitive flexibility, i.e. the ability to rapidly adapt to a change in the task contingency, in the context of abstract reasoning [8]. Early studies utilizing and developing scoring conventions for the test [35] focused on perseverative errors. However, it has since become clear that the test does not engage just a single cognitive process for task set switching; rather, it relies on a variety of cognitive functions throughout the test including working memory, attention, decision making, inhibitory control and reasoning [21, 36, 37, 38, 39]. For example, the “failure to maintain set” error occurs when a subject applies an incorrect rule after they have learned the correct rule and is associated with an error in working memory.

This has inspired systematic studies on WCST performance with two related goals. First, research has focused on an accurate characterization of rule-learning strategies and/or the cognitive processes that support their underlying computations [17, 20, 21]. Here we developed a relatively hypothesis-free approach to identify the rule-learning strategy in humans and monkeys based on hidden behavioral states. The best-fit models for both species ascribe these hidden states to varying levels of attention to individual task-relevant visual features (Fig. 3a). These results are consistent with the conclusions of earlier studies that humans contend with the “curse-of-dimensionality” which is inherent in the WCST with selective attention towards individual features during exploration [17, 20, 21]. Our findings clarify these results by showing that in the high-dimensional version of the task (twelve instead of three possible rules) both humans and monkeys must further contend with a tradeoff between computational complexity and information efficiency while exploring for the rule, and they do so by selectively attending to a few, but not all, features at a time (Fig. 4e).

Our approach differs from these earlier studies in that it does not postulate a specific learning algorithm [17, 20]. Rather, it discovers the decision process that determines the rule-learning strategy. In doing so, it illustrates important differences between human/monkey rule learning strategies on the WCST and the commonly observed winstay lose-shift learning strategy (Fig. 3b). For example, a key function of the preferred state, which is not part of the win-stay lose-shift strategy, is to support the simultaneous exploration of multiple features at a time over many trials. Moreover, this state is also associated with inference-like computations that support a computationally efficient strategy of narrowing down the rule by eliminating other candidates using unambiguous negative feedback. Indeed, a reinforcement-learning inspired description of rule learning in the WCST by Bishara et al. [20] was parameterized to update the value of unchosen options facilitating inference-like computations. However, the prevalence of the behavior itself was not reported by the authors.

The second goal, with stronger clinical implications, is a comparison of error types in healthy and diseased populations towards identifying more accurate behavioral markers for different types of neuropsychiatric disorders and for dysfunction or lesions of different brain regions [12, 19, 20, 21, 38, 39, 40, 41, 42, 43]. For example, a detailed analysis of non-perseverative errors led to the differentiation between “efficient” errors that are expected to occur during hypothesis testing from “random” errors that reflect a failure in maintaining the cognitive set, and demonstrated a prevalence of the latter in patients with frontal lobe pathology [19]. In support of this goal, we have developed a learning-stage categorization method that delineates learning stages by the features under exploration and their relationship to the rule (Fig. 5a). Intuitively, this approach tracks how far along a subject is from learning the rule and reflects this in the reward rates across categories (Fig. 5c, bottom). Furthermore, the categories are mutually exclusive and exhaustive, allowing us to precisely ascribe differences in learning performance between subjects or even between rule-blocks for the same subject to differences in individual categories (Fig. 5a; Supplementary Fig. 7).

Importantly, the approach identifies various known classes and sub-classes of error types, but also newer ones that may prove useful in future investigations of behavioral markers for neuropsychiatric disorder and cognitive impairment. It distinguishes perseverative errors (made during the perseveration category) from non-perseverative ones. Consistent with earlier work [19], it further sub-categorizes the latter into random and efficient errors. It identifies two forms of random errors: one occurs during rule search (before the rule preferred or exploitation categories) when subjects occasionally choose none of the features they are currently exploring (Fig. 4d); the other occurs after they have found the rule and while they are demonstrating this (Fig. 6b). However, it remains unclear if either of these random errors are a feature of cognitive flexibility and result from random exploration, or a bug caused by the failure to maintain the attention set in working memory. Indirect evidence in humans has been found in favor of the latter interpretation [44]. If in fact it is a result of higher distractability in monkeys, monkey performance may be improved by imposing stronger controls on potential environmental distractors [45]. Efficient errors arise instead when subjects test hypotheses regarding rule identity and occur during the random search and non-rule exploration categories. However, most errors during the rule-favored exploration category are repeated despite unambiguous (direct or indirect) negative feedback (Fig. 7). These “disambiguation” errors are neither random nor efficient but arise from a deficit in disambiguating the rule feature that is under exploration from a simultaneously explored non-rule feature. This newly identified error type bears further exploration in patient populations.

The higher incidence of these error types in monkeys may have more to do with how they learn rather than some fundamental cognitive constrains — since they cannot receive a rich verbal description of the task’s structure as humans do and must learn about it via trial-and-error, it is possible that monkeys misinterpret uncued rule switches as stochasticity in the environment resulting in a maladaptive strategy. Nevetheless, it is noteworthy that many of the errors we have identified as contributing to deficits in monkey rule-learning performance have also been implicated in the poor performance of humans with cognitive impairment. A higher incidence of perseverative errors in patients with prefrontal cortex pathology was first reported in a landmark study by Milner [10]. Random errors, while rare in healthy humans, are more frequently observed in patients with frontal lobe dysfunction [19]), similar to the present study finding in monkeys. Moreover, poor sensitivity to negative feedback, which underlies perseverative and disambiguation errors in monkeys, is more pervasive in patients with schizophrenia and substance abuse as well [20, 21].

Ultimately, these questions can be resolved by inter-species comparisons of neural data. For example, perseverative and random errors have distinct neural signatures in humans [46]; are similar signatures observed in monkeys? More generally, our work produces several testable neural hypotheses in both species. First, does the neural representation of the current rule persist during and across rule exploitation trials (i.e. after its identity has been learned) [47, 48, 49, 50]? Second, since the set of explored features must be maintained across several trials, are they represented by neural activity during and across trials? Our model indicates that this attention set is typically small (Fig. 4e) and longer bouts of exploring multiple features simultaneously (Fig. 7b) requires a choice alternation between these features. This drives the need to maintain the explored features in working memory, particularly to support recall of one of them after it is not chosen on one or more previous trials. Third, are the distinct error types (perseverative, random, efficient, disambiguation) differentially represented in the brain? Error coding neurons have been reported in the prefrontal cortex of monkeys performing a WCST analogue [50, 51]. Moreover, perseverative, random and disambiguation errors signal the need to disengage from the previous rule, address a working memory error and remove a non-rule feature from the attention set, respectively. This difference in their function raises the possibility that they are represented differently, either eliciting stronger responses in different brain regions or eliciting differential responses in the same region [46]. Fourth, is the strength of these error signals or their modulation of the attention set representation [50] larger on trials when they serve their function? For example, are perseverative error signals stronger on trials after which perserveration halts compared to those that are followed by continued perseveration? A potential reason for inter-species performance differences may be found in these analyses: what inter-species neurocognitive differences explain the relative prevalence of perseverative, random and disambiguation errors in monkeys compared to humans, and are they also observed in humans with cognitive impairment?

There exist several avenues to clarify and improve upon our modeling approach. A key difference between earlier models and ours is our assumption that each feature is associated with discrete states, which our model relates to feature-based attentional states. In contrast, Bayesian and reinforcement learning approaches posit that subjects reason about features by assigning continuous-valued functions such as belief [17] and value [18, 20] to them, respectively. In future work, we will test whether a model with continuous-value states provide a better fit to the behavior of the two species, which may offer a different interpretation of the latent variable used by them to reason about features. Our model has also been simplified to keep subsequent analysis tractable — it does not explicitly account for interactions between features. This had the unintended consequence of discovering the “phantom” avoid state. Future improvements to our model will carefully incorporate such interactions explicitly, while retaining high interpretability.

In conclusion, we have applied a hypothesis-free state-characterization method to identify and compare the rule-learning strategy on the Wisconsin Card Sorting Test in humans and monkeys. The hidden attentional states and state transitions inferred by the model facilitated the determination of the decision process underlying this strategy as well as the various stages of rapid rule learning. The inferred states perfectly (substantively) explain human (monkey) choice behavior (Fig. 2c). Our overall approach reveals differences in cognitive strategy between the two species and isolates the identity and relative contribution of various error types to the performance difference between the two species. It shows that random exploration or distraction and poorer sensitivity to negative feedback underlies a higher incidence of these error types in monkeys thus leading to their under-performance. The high fidelity demonstrated by the model in inferring hidden attentional and decision states holds promise in advancing the search for more accurate behavioral markers of various types of cognitive dysfunction and in motivating targeted analyses to determine and compare the neural correlates of the various cognitive processes engaged by the WCST.

## Acknowledgements

We thank S.W. Linderman for generously sharing his code and advice on fitting HMM-GLM models to data; I.R. Stone, J. Ferre and E.Y. Walker for fruitful discussions. This work was supported by the National Institute of Health U-19 program grant no. 5U19NS107609-03, R01 grant no. R01MH062349 and the Office of Naval Research grant no. N00014-17-1-2041.

## Data and Code Availability

All training and analysis codes will be available at publication on GitHub (https://github.com/xjwanglab). We will also provide data files in Python readable formats for further analyses. Pre-trained models will be stored in a Google Drive folder with its link provided on the same GitHub repository.

## Methods

### Task Description

Human and monkey subjects were tested on an analogue of the Wisconsin Card Sorting Test (WCST). On each trial, they were simultaneously presented with an array of four objects on a computer screen. Each object was comprised of a stimulus feature from each of three stimulus dimensions: color, pattern, and shape (Fig. 1a), e.g., a blue polka-dotted triangle. For each trial, these four objects were chosen from a pool of 64 unique objects, each containing a possible combination of individual features from each of the three dimensions, such that there was no feature overlap between them. Accordingly, for each possible array, all four features of each dimension appeared on the screen, but the combination of features represented by each individual object varied across trials. Within a single rule-learning block of trials, one color, texture, or shape was designated as the target, resulting in 12 possible rules. The identity of this rule was not cued, but had to be learned by trial and error, based on the feedback received at the end of each trial. Upon meeting a rule-learning criteria for the current rule, the rule feature changed on the next trial in an uncued manner, initiating a new rule-learning block. This rule shift could be either intradimensional, where the dimension of the new rule feature matched that of the previous rule feature (e.g., changing from triangle to square), or extradimensional, where the dimensions of the old and new rule features did not match (e.g., changing from triangle to yellow);

### Monkeys

All procedures were carried out in accordance with the National Institutes of Health guidelines and were approved by the University of Washington Institutional Animal Care and Use Committee. Subjects were four adult female rhesus monkeys (Macaca mulatta) with mean age 12.5 *±* 2.5 years and mean weight 7.5 *±* 0.6 kg at the start of the experiment. Prior to testing, a titanium post for holding the head was surgically affixed to each monkey. During testing, each monkey was head-fixed in a dimly illuminated room and positioned 60 cm away from a 19-inch CRT monitor with a screen refresh rate of 120 Hz noninterlaced. The monitor had a resolution of 800 *×* 600 pixels, subtending 33 degrees by 25 degrees of visual angle (dva). Eye movements were recorded using a noninvasive infrared eye-tracking system (EyeLink 1000 Plus, SR Research). Stimuli were presented using experimental control software (NIMH Cortex or NIMH MonkeyLogic). Calibration of the infrared eye tracking system was accomplished using a nine-point manual calibration task.

Following the calibration task, the monkey was tested on the WCST analogue. The monkey initiated each trial by fixating a white cross (0.5°) at the center of the computer screen. Following 500 ms of successful fixation, the cross disappeared and was replaced by an array of four objects. During the self-paced decision epoch that followed, the monkey was free to explore the array of objects; her response was defined as maintaining her gaze within a 9° *×* 9° window centered on the object for 800 ms. The monkey received a food slurry reward over a 1.4-second duration for selecting the object that contained the rule feature. A time-out period (either 1-second or 5-seconds) occurred on trials where the monkey did not choose the object containing the rule feature or where she did not make a choice within 4 seconds. The feedback period was immediately followed by a 400 ms or 1 s inter-trial interval. We classified a rule as learned either when the monkey made eight consecutive correct responses or when she made 16 correct responses in 20 trials or fewer.

The type of rule shift that followed (intradimensional or extradimensional) was determined pseudo-randomly to occur with equal probability. A block consisted of all the trials from the initial rule shift to the final trial of criterion performance. We analyzed a total of 1305 blocks in 81 recording sessions from monkey B, 872 blocks in 29 recording sessions from monkey C, 805 blocks in 29 recording sessions from monkey S, and 224 blocks in 13 recording sessions from monkey T. Only completed blocks were included in the analysis.

### Humans

The studies involving human participants were reviewed and approved by the Institutional Review Boards of University of California, Berkeley. All participants provided their written informed consent to participate in this study and received a small compensation. These subjects were four adult males and one adult female with mean age 26.4 *±* 4.1 years. Subjects were brought into a room where they sat and completed a computer-adapted version of WCST analogue on a recording laptop after receiving the following instructions: “In this experiment, you will see 4 cards on each trial. Each card has 3 unique features (color, shape and texture). No feature is shown on more than one card, so you will see 12 different features on each trial (4 colors, 4 shapes, 4 textures). The card containing the correct feature (1 out of 12 possible) will be correct choice. The correct feature might change during the task. The answer is given by pressing one of the four arrow keys that corresponds with the selected card position on the screen (up, down, left or right). You have 4 sec to provide the answer, or the trial times out. The task goes on for 200 trials or about 15 minutes.”

Individual trials consisted of the following epochs: cross fixation (black cross displayed in the center of the screen on a gray background for 300 ms), choice (four objects displayed on the screen at locations corresponding toup, down, left or right positions, for up to 4000 ms), feedback (‘correct’ or ‘incorrect’ feedback message displayed for 1500 ms) and inter-trial interval (ITI, gray screen for 1000 ms). Subjects indicated their choice by pressing the arrow key on the laptop keyboard, corresponding to the chosen object’s position on the screen. If the choice was not indicated within the 4000 ms, the trial was considered timed-out. After reaching the learning criteria, defined as 5 consecutive correct trials or 8 correct out of the last 10 trials, the rule was switched and a new rule-learning block began. The new rule was randomly determined. Each participant completed five task sessions (300 trials/sessions for a total of 1500 trials). This spanned between 107 and 138 blocks across the five subjects.

### Win-Stay Lose-Shift (WSLS) Agent

The task structure (rule selection, learning criteria) for the WSLS agent was identical to that of the humans, except for the trial structure – the agent’s algorithm determined its choice immediately upon stimulus presentation. The algorithm (Supplementary Fig. 3d, left) always maintained a single feature in the persist state and deterministically chose the object with that feature at each trial. Positive feedback maintained the feature in the persist state. Negative feedback demoted it to the avoid state and promoted, a randomly selected feature from among the 11 others that were in the avoid state, to the persist state. The agent completed 500 rule blocks.

## Input-Output Hidden Markov Model - Generalized Linear Model (IOHMM-GLM) for the prediction of feature choices

### Model Design

The four objects presented during a trial consist of twelve visual features, *f* ∈ *{*1, …, 12*}*. In support of feature-based mental representations, the model predicts the choice of each feature *f* at the next trial *t*. This choice is represented by 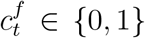, where 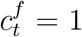 indicates the *f* was part of the chosen object, and 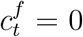 indicates it was not.

Either choice can result in a reward or timeout for the trial, given by *r*_*t*_ ∈ *{*0, 1*}*. The choice-outcome history of *f* given the past ℓ trials is denoted *h*^*f*^ ∈ *{*1, …, 2^2ℓ^ *}*. We refer to ℓ as the *lag* and it is a hyperparameter of the model. The value of *h*^*f*^ at trial *t* is given by the binary vector (*r*_*t*−1_, *c*_*t*−1_, …, *r*_*t*−ℓ_, *c*_*t*−ℓ_) of size 2ℓ. Therefore, it can take on 2^2ℓ^ possible values. In all our analyses, we choose a lag 1 (ℓ = 1) model for further analysis. Such a model depends on a choice-outcome history that takes on one of four possible values at trial *t*: (*r*_*t*−1_ = 0, *c*_*t*−1_ = 0), (*r*_*t*−1_ = 1, *c*_*t*−1_ = 0), (*r*_*t*−1_ = 0, *c*_*t*−1_ = 1) or (*r*_*t*−1_ = 1, *c*_*t*−1_ = 1) which we refer to as *NC*^−^, *NC*^+^, *C*^−^ and *C*^+^ respectively.

The transformation of the choice-outcome history into a choice at trial *t* is mediated by discrete hidden states *s*^*f*^ ∈ *{*1, …, *K}* that determine the parameters of the transformation. The maximum number of states *K* is a second model hyperparameter. The transformation is modeled as a Bernoulli GLM:

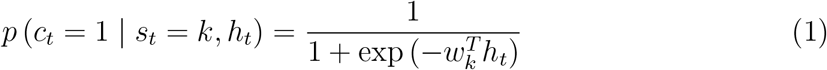

where the parameters 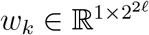 are determined by the state *s*_*t*_ = *k*. We denote the set of parameters across all *K* states as 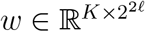.

Transitions between states also depend on the choice-outcome history and are modeled by multinomial logistic regression:

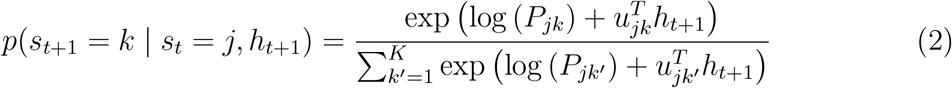

where the parameters 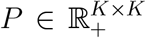 and 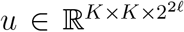 represent the bias or baseline transition probability and history weights. This model design is schematized in Figure 2a.

Finally, the probability distribution of initial states *π*, is a model parameter that specifies the state at the first trial of a session.

### Model Fitting

We fit the parameter values for the choice GLM weights *w*, the baseline transition probability *P*, the transition GLM weights *u* and the initial state distribution *π* to the choices of each subject. To avoid over-fitting, the parameter values were shared across all features. In other words, all parameter values were the same for all 12 features. The likelihood of the data under a model is its probability subject to the model’s parameters and inputs *p*(*c*_1..*T*_ | *w, P, u, π, h*_1..*T*_), where *T* is the number of trials in the session. It is expressed in terms of these parameters as:

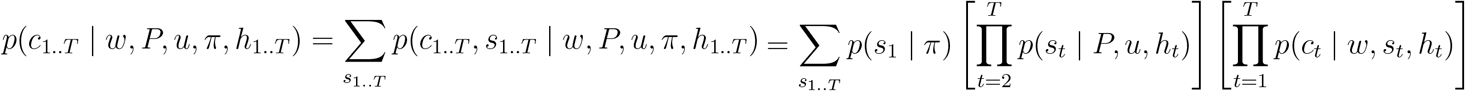

where the last two terms are given by equations 2 and 1 respectively.

The model parameters were fit by minimizing −*log* [*p*(*c*_1..*T*_ | *w, P, u, π, h*_1..*T*_)], i.e. the negative log-likelihood of the data, via gradient descent with the ADAM optimizer. The choice GLM weights for all *k* states were initialized to a single 2^2ℓ^ -dimensional vector drawn from a standard normal distribution. The baseline transition probability was initialized to the sum of a diagonal matrix with value 0.9*I* where *I* is the identity matrix, and a random matrix with elements drawn from a uniform distribution in the interval [0, 0.05). The larger diagonal values enforce “stickiness” that bias transitions back into a state. The transition GLM weights were initialized to zero, and the initial state distribution was initialized to 1*/K* for each state *k*. For each subject and each pair of hyperparameters (ℓ, *K*), the parameters were optimized over 10000 iterations with 5-fold cross validation (Fig. 2b).

The best fit model was sought for each human (monkey) subject and hyperparameter setting across 10 (5) independent parameter initializations. Figure 2b shows the mean negative log-likelihood taken over all initializations and cross-validation folds. The best-fit model for each human (monkey) subject was selected for further analysis from these 50 (25) models at hyperparameter values ℓ = 1 and *K* = 4. Although, we found that a majority of these models produced very similar choice and transition probabilities. However, fits to the WSLS agent varied much more. Since negative feedback immediately demoted features from the persist to avoid state, exploration of non-rule features typically lasted 1-2 trials. This likely makes it harder for the model to identify exploration and introduces more variability across fits.

Once the best-fit model is identified, the most likely sequence of states, *s*^*^, for each subject, session and feature in determined by the Viterbi algorithm [1] (Fig. 2d). For each trial *t* and feature *f*, the algorithm performs a forward pass across all past trials and a backward pass across all future trials to determine the most likely state of *f* at trial *t* that best explains past, present and future history-dependent choices under the constraints of the model’s parameters and the choice and transition probabilities they yield. Supplementary Figure 2b) shows the cumulative distribution of the posterior probabilities 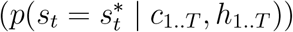 of these state estimates calculated for the Viterbi algorithm.

All model fits and the most-likely state determination was performed with the State Space Model (SSM) python package [2].

### Model Extension for the Prediction of Object Choices

We extended the feature choice prediction model described in the previous section to predict object choices at each trial *t*. Given the predicted choice probability 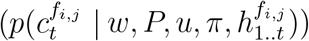 for each feature *fi,j, i* ∈ *{*1, …, 3*}* in an object *o*_*j*_, *j* ∈ *{*1, …, 4*}* presented at trial *t*, the model predicts the object chosen at *t*. This transformation of predicted feature choice probabilities *p*(*f*_*i,j*_) into object choice probabilities *p*(*o*_*j*_ | *p*(*f*)) is modeled by multinomial logistic regression:

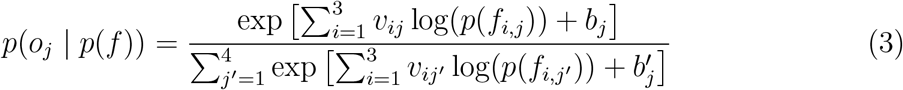

where the parameters *v* ∈ ℝ^3×4^ and *b* ∈ ℝ^1×4^ represents the feature choice probability weights and biases in selecting each object, respectively. These values were fit to the choices of each subject by minimizing the cross-entropy loss 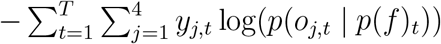 where *y*_*j,t*_ ∈ *{*0, 1*}* indicates whether object *o*_*j,t*_ was chosen on trial *t*. Model fitting was performed via stochastic gradient descent with the ADAM optimizer implemented by the Pytorch python package [3]. The parameter values for *v* and *b* were initialized from a uniform distribution in the interval 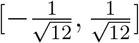 and optimized until convergence with a maximum of 100000 iterations. Cross validation was performed with the same training and test sets used while training the feature choice prediction models (Supplementary Fig. 2a).

The accuracy of the object choice prediction model based on the best-fit feature choice prediction model with 4 states and lag 1 is shown in Figure 2c, left. We also fit a model to determine the chosen object in a similar fashion using the feature choice probabilities based on their most-likely state estimates 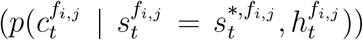 instead. The accuracy of this model is shown in Figure 2c, right.

### Model Analysis

The probability distribution of histories in each state (Supplementary Fig. 4b) is:

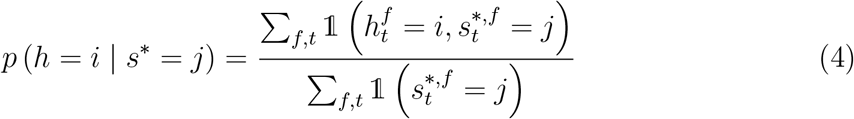

where 𝟙 is the indicator function and ∑ _*f,t*_ is a sum over features and trials. The state and history dependent choice probability (Supplementary Fig. 4a) can be directly calculated from the model’s parameters (Eqn. 1) or empirically as:

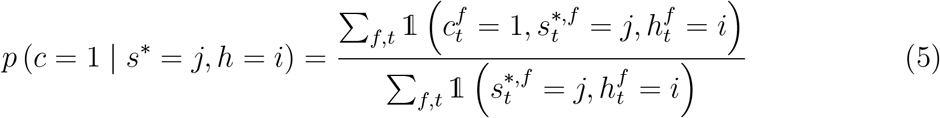

The choice probability of a feature in each state (Fig. 3a) can be computed by utilizing equations (4) and (5) or 1.

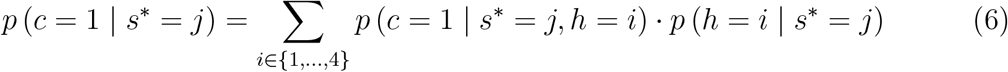

Similarly, the state transition probabilities (Supplementary Fig. 5) can be directly calculated from the model’s parameters (Eqn. 2) or empirically as:

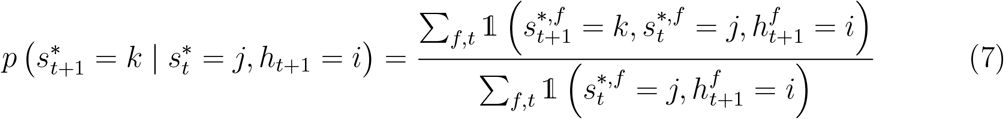

We approximated the decision process in each species (Fig. 3b) from the state transition probability, and the “reverse” state transition probability 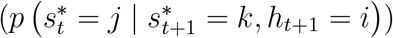. The latter helps in conditions where transitions into a state are typically rare, such as transitions from the random/avoid state into the preferred state. This quantity (Supplementary Fig. 6) is calculated empirically as:

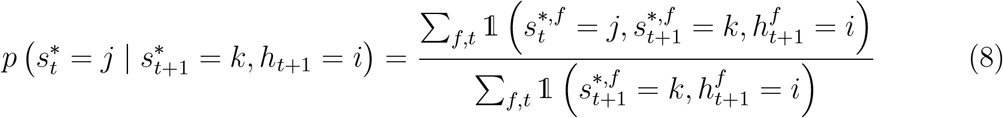

### Trial Categorization

Trials were categorized based on the identity of the rule feature and the most-likely state estimates for all 12 features as in Figure. 5a. Since each trial is always designated to one and only one category, the trial categories are mutually-exclusive and exhaustive. This facilitates a precise decomposition of the length of each rule block into the number of trials spent in each category (Fig. 5c). Moreover, since the categories are mutually exclusive, we can explain summary statistics (mean and variance) of the block length for each subject in terms of statistics of their category lengths (Supplementary Fig. 7):

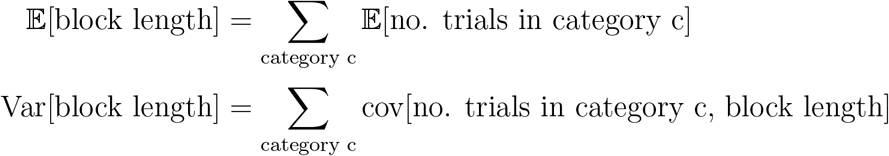

### Inter-species Comparison of Category Lengths

In Figure 7, the higher probability of continued exploration of non-rule features by monkeys during the rule-favored exploration category is attributed to poor (direct and indirect) negative feedback sensitivity (Fig. 7c-d). In addition, we attribute the higher probability of continued exploration in monkeys during rule-favored exploration trials compared to non-rule exploration trials, to a higher prevalence of direct positive feedback during rule-favored exploration trials (Fig. 7f).

These determinations were made based on the following decomposition:

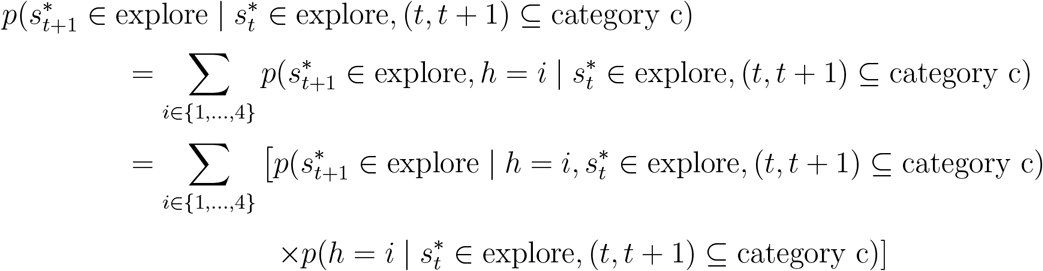

The joint probability above is shown in Figure 7f, left and in Supplementary Fig. 8), and quantities resulting from its decomposition below are shown in Figure 7f, middle-right.

## Supplementary Tables and Figures

**Supplementary Table 1:** Inter-species trial structure differences.

|  | Fixation | Decision | Response | Feedback | ITI |
| --- | --- | --- | --- | --- | --- |
| Monkeys | 500 ms | $\leq 4$ s | Fixate<br>(800 ms) | 1.4 s (reward)<br>1 s / 5 s (timeout) | 400 ms /<br>1 s |
| Humans | 300 ms | $\leq 4$ s | Manual | 1.5 s<br>(text on screen) | 1 s |

**Supplementary Table 2:** Inter-species learning criteria differences.

|  | Learning Criteria:<br>Continuous | Learning Criteria:<br>Proportion |
| --- | --- | --- |
| Monkeys | 8 correct trials | 16/20 correct trials |
| Humans | 5 correct trials | 8/10 correct trials |

**Supplementary Figure 1:**
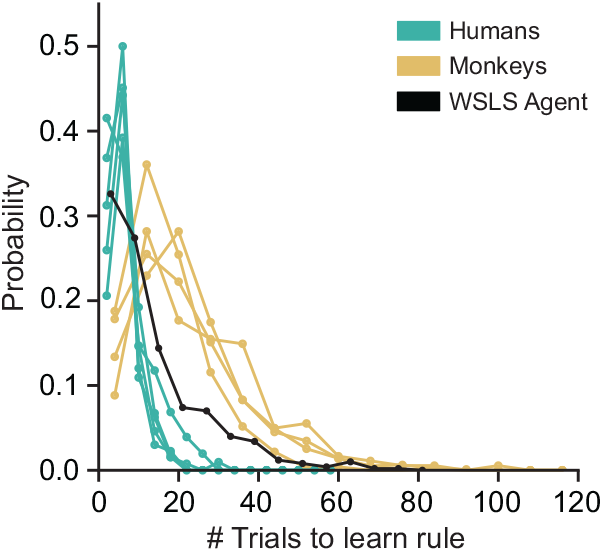
Monkey learning is slower than humans after correcting for learning criteria differences. Distribution of trials-to-learning-criteria in 4 monkey (brown) and 5 human (green) subjects. The less stringent learning criteria used in human subjects was applied to the monkey choices and outcomes to revise their trials-to-learning-criteria on each rule block. Yet, monkeys are over 3 times slower than humans on average. Humans are also faster than a simulated agent using the Win-Stay Lose-Shift strategy to learn WCST rules (black).

**Supplementary Figure 2:**
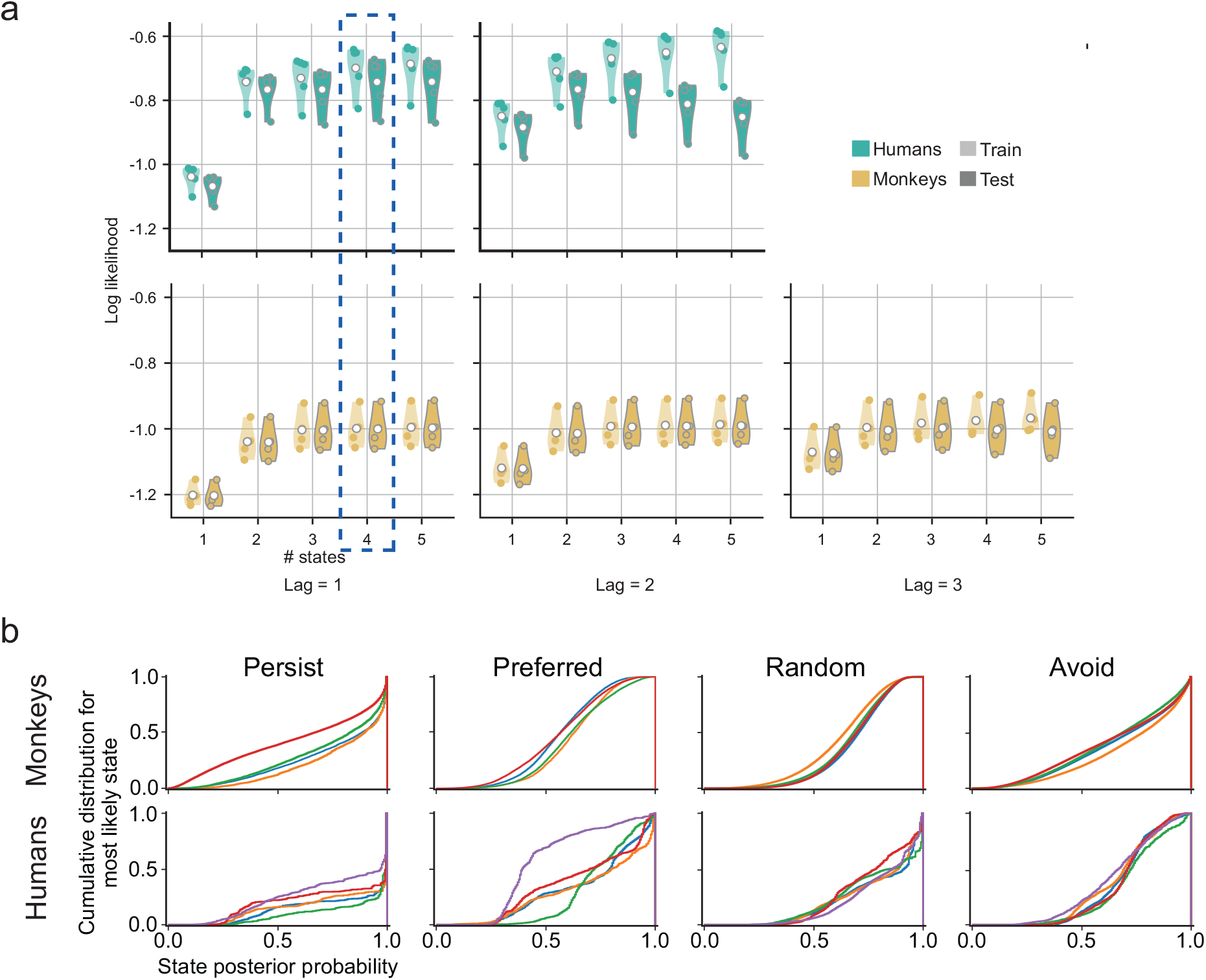
Goodness of fit of a model extension to predict chosen objects is consistent with the underlying feature prediction model. **a**. Log-likelihood of a model extension that predicts the chosen object from feature choice probabilities. Log-likelihoods are show separately on training and test datasets for each human (green) and monkey (brown) subject for extensions of feature choice models with varying numbers of states and that use choice outcomes from varying numbers of previous trials (lag). Each point represents the mean results over a 5-fold cross-validation and over 5 (monkey) or 10 (human) different initializations. **b**. Cumulative density of the posterior probabilities for most-likely state estimates by the model. Densities are presented separately for the 4 states and for each monkey (top) and human (bottom) subject.

**Supplementary Figure 3:**
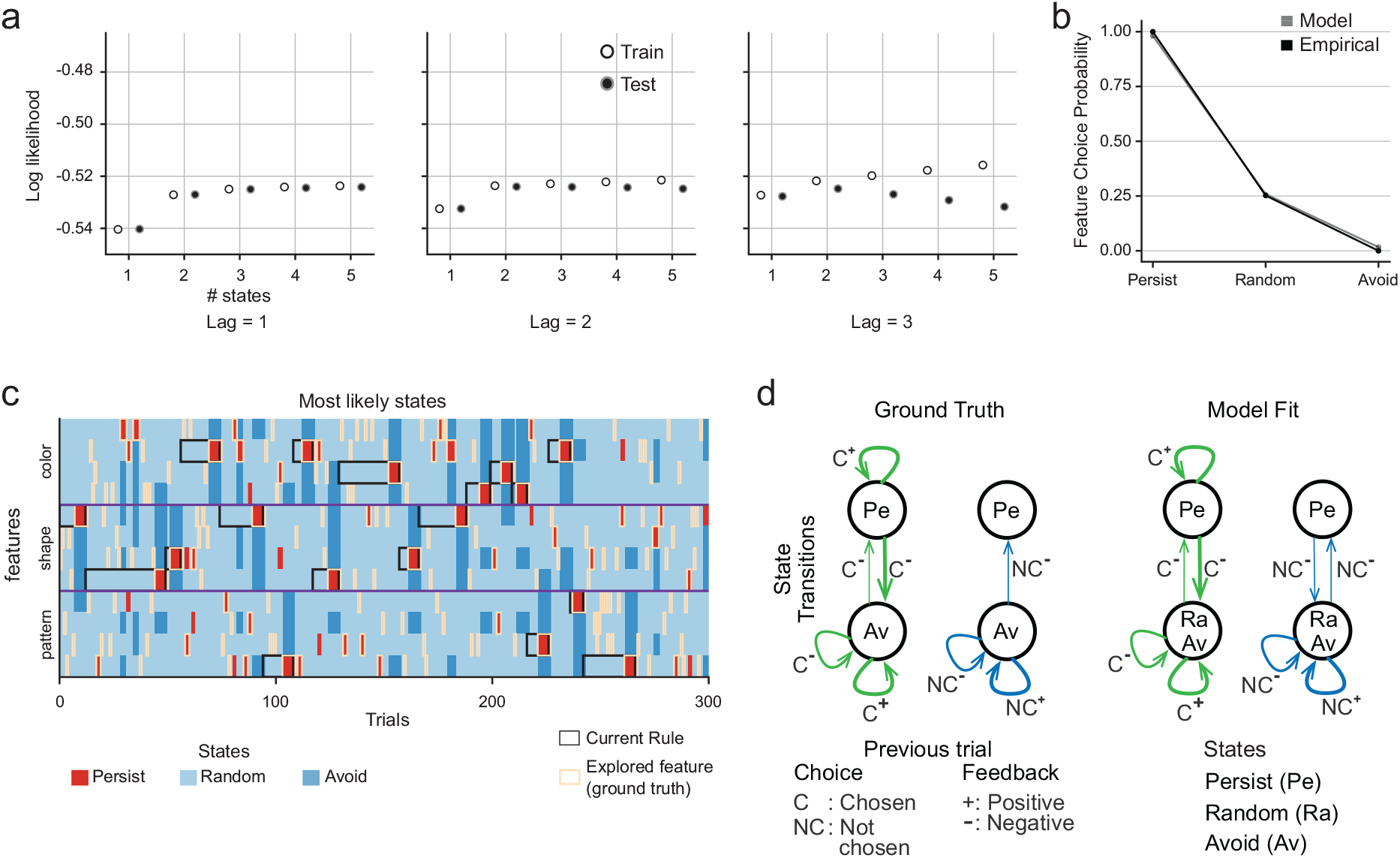
Model recovers strategy of a simulated Win-Stay Lose-Shift agent. **a**. Model fit log-likelihoods for training and test datasets generated by a simulated agent using the Win-Stay Lose-Shift strategy on the WCST. The lag and number of states were parametrically varied across models. Each point represents the mean over a 5-fold cross-validation and over 10 different model initializations. **b**. Probability of choosing a feature associated with each state of the best-fit 3-state, lag-1 model. Plot shows probabilities generated by model parameters (gray) and measured empirically based on the most-likely state estimates (black). **c**. Most-likely states estimated by the model for 300 example trials. The rule on each block is outlined in black, and the ground-truth feature under agent exploration on each trial is outlined light brown. 57.6% of all ground-truth features under exploration are accurately assigned the persist state (choice probability = 1) by the model. **d**. Decision process for the Win-Stay Lose-Shift learning strategy employed by the agent (ground-truth; left) and fit to the agent’s choices (right). The ground-truth process is modified from the one in Figure 1c to account for the composition of the chosen object by 3 features - the feature under exploration and two features that are extra-dimensional to it. Features in these other two dimensions are chosen even though they are not under exploration. The model differentiates these randomly chosen features from those that are intra-dimensional to the explored feature and completely avoided. The fit recovers the underlying decision process.

**Supplementary Figure 4:**
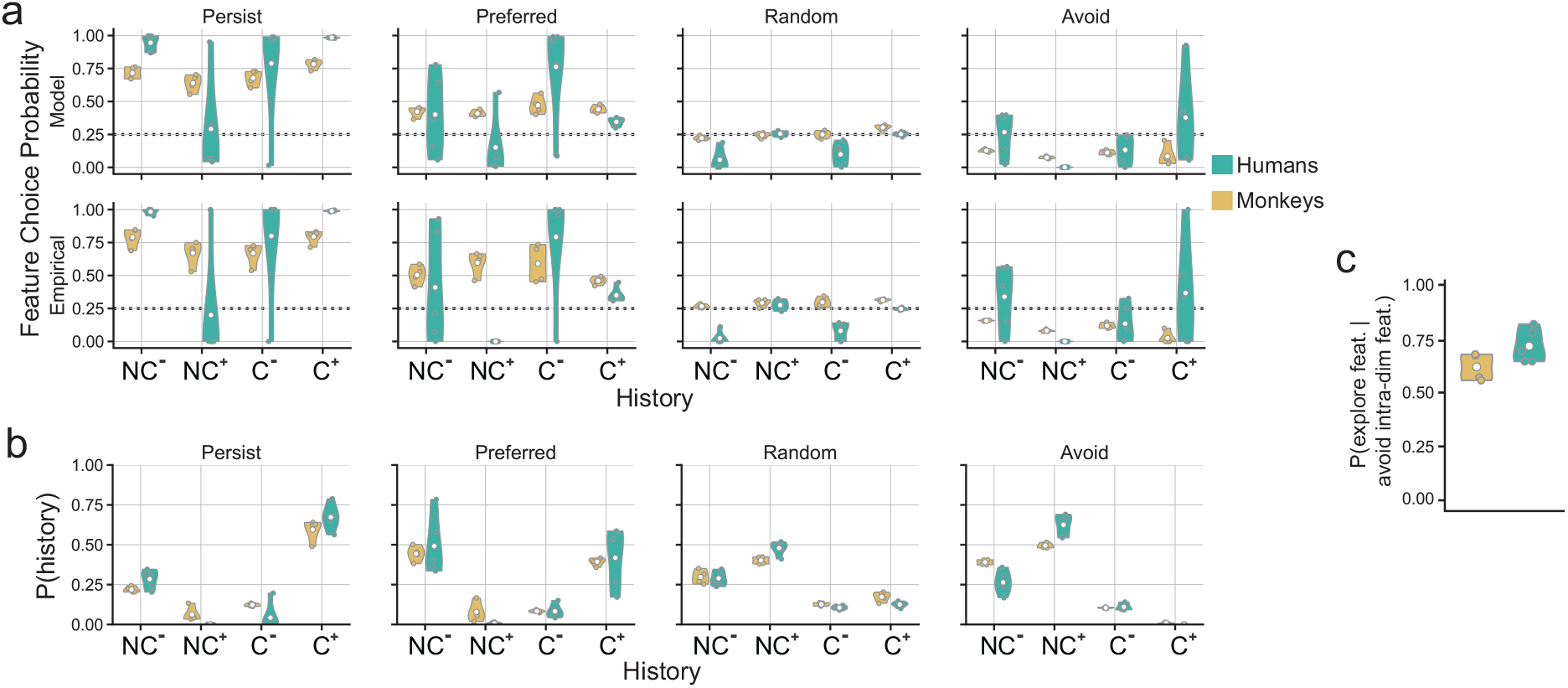
State- and history-dependent statistics. **a**. Feature choice probability given the feature’s state and history in human (green) and monkey (brown) subjects. Values computed directly from model’s parameters (above) are consistent with empirical measurements based on best-fit state estimates (below). **b**. Probability distribution of histories given the state estimate on the subsequent trial. **c**. Probability that a feature is associated with the preferred or persist state given one of its intra-dimensional counterparts is associated with the avoid state.

**Supplementary Figure 5:**
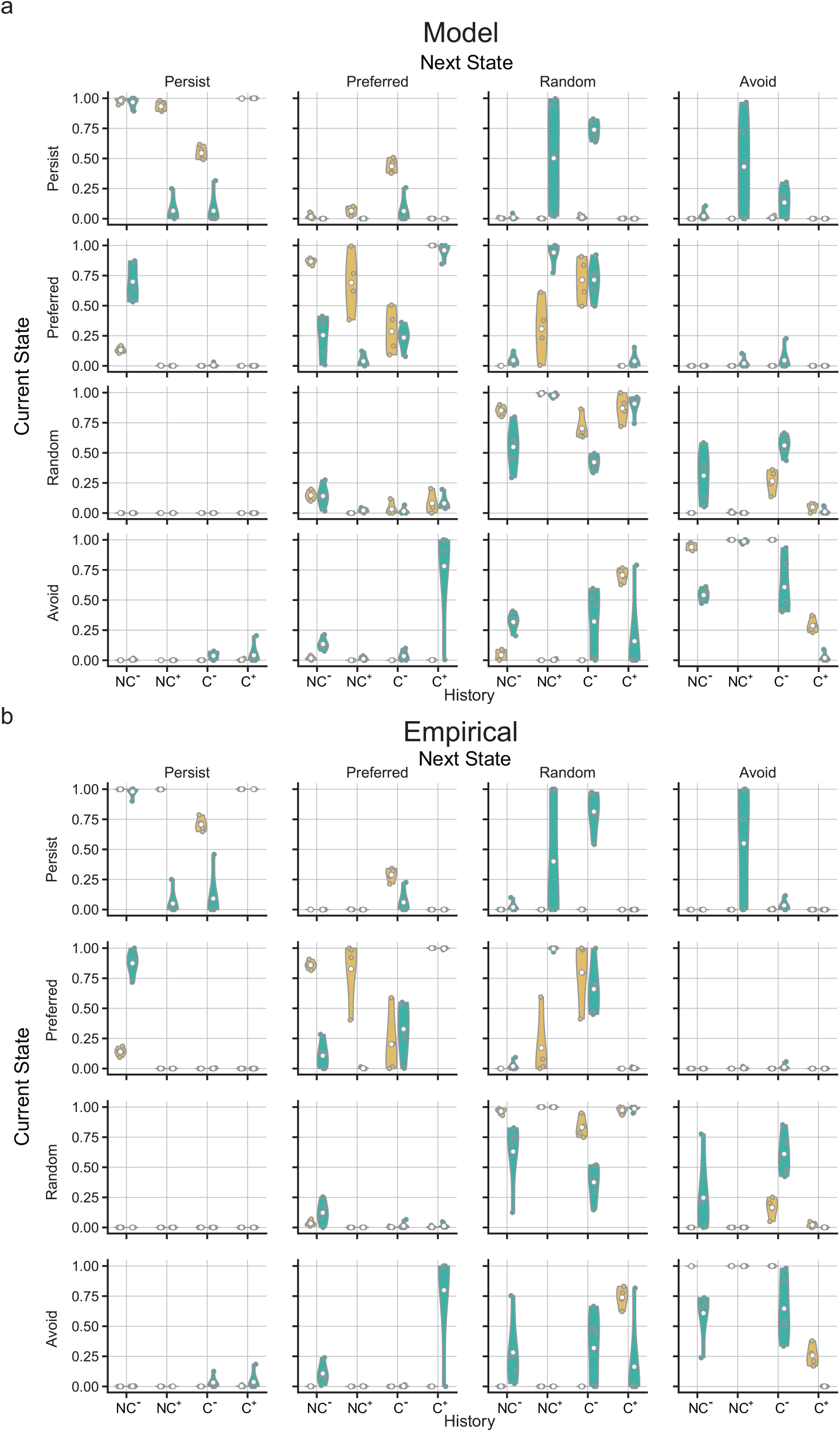
State transition probabilities. **a-b**. Probability of a feature’s state transitions given the state it was associated with on the previous trial and its choice-outcome history in human (green) and monkey (brown) subjects. Values computed directly from model’s parameters (a) are consistent with empirical measurements based on best-fit state estimates (b).

**Supplementary Figure 6:**
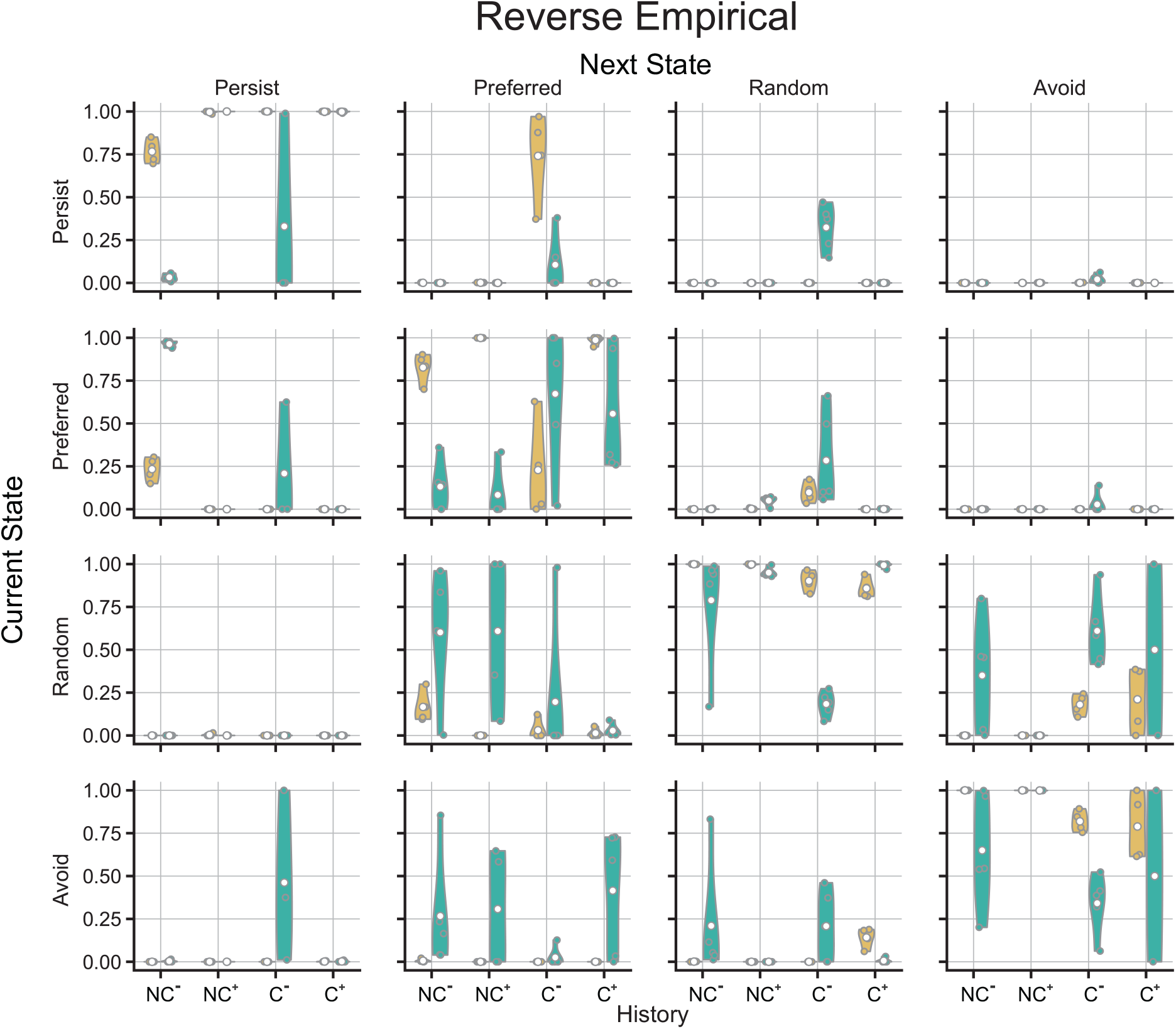
‘Reverse’ state transition probabilities. Empirically measured probability of a feature’s state transitions given the state it is associated with on the next trial and its choice-outcome history in human (green) and monkey (brown) subjects.

**Supplementary Figure 7:**
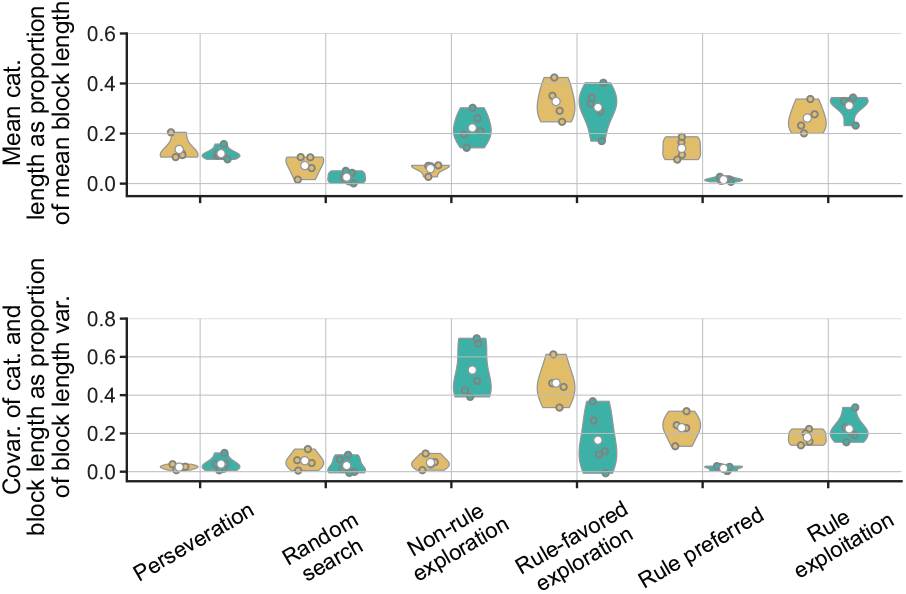
Trial categories fully determine block length statistics. Mean category length as a fraction of mean block length (top) for human (green) and monkey (brown) subjects. Covariance of category length with block length as a fraction of the variance in block length (bottom). Since the categories are mutually exclusive and span all trials in a block, the sum over categories of each of these two fractions is 1. This allows an assessment of each category’s contribution to the mean (top) and variance (bottom) of the block length.

**Supplementary Figure 8:**
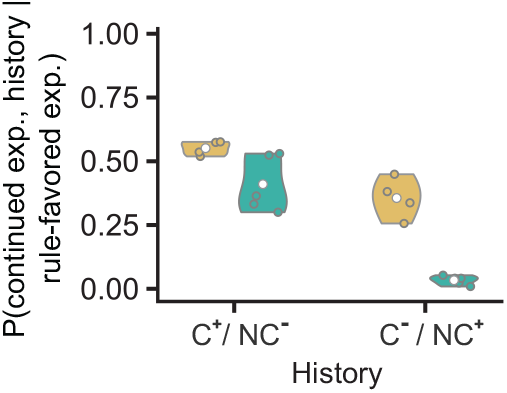
**Joint probability of a non-rule feature transitioning back into an exploration state and receiving direct/indirect positive (**C^+^/NC^−^**) or negative (**C^−^/NC^+^**) feedback during rule-favored exploration trials in human (green) and monkey (brown) subjects**.

